# m^6^A modification of U6 snRNA modulates usage of two major classes of pre-mRNA 5’ splice site

**DOI:** 10.1101/2022.04.05.487178

**Authors:** Matthew T Parker, Beth K Soanes, Jelena Kusakina, Antoine Larrieu, Katarzyna Knop, Nisha Joy, Friedrich Breidenbach, Anna V Sherwood, Geoffrey J Barton, Sebastian M Fica, Brendan Davies, Gordon G Simpson

**Affiliations:** School of Life Sciences, University of Dundee, Dow Street, Dundee, DD1 5EH, UK; Centre for Plant Sciences, School of Biology, Faculty of Biological Sciences, University of Leeds, Leeds LS2 9JT, UK; Department of Biochemistry, University of Oxford, South Parks Road, Oxford OX1 3QU, UK; Cell & Molecular Sciences, James Hutton Institute, Invergowrie, DD2 5DA, UK

## Abstract

Alternative splicing of messenger RNAs is associated with the evolution of developmentally complex eukaryotes. Splicing is mediated by the spliceosome, and docking of the pre-mRNA 5’ splice site into the spliceosome active site depends upon pairing with the conserved ACAGA sequence of U6 snRNA. In some species, including humans, the central adenosine of the AC<u>A</u>GA box is modified by *N^6^* methylation, but the role of this m^6^A modification is poorly understood. Here we show that m^6^A modified U6 snRNA determines the accuracy and efficiency of splicing. We reveal that the conserved methyltransferase, FIO1, is required for Arabidopsis U6 snRNA m^6^A modification. Arabidopsis *fio1* mutants show disrupted patterns of splicing that can be explained by the sequence composition of 5’ splice sites and cooperative roles for U5 and U6 snRNA in splice site selection. U6 snRNA m^6^A influences 3’ splice site usage and reinforces splicing fidelity at elevated temperature. We generalise these findings to reveal two major classes of 5’ splice site in diverse eukaryotes, which display anti-correlated interaction potential with U5 snRNA loop 1 and the U6 snRNA AC<u>A</u>GA box. We conclude that U6 snRNA m^6^A modification contributes to the selection of degenerate 5’ splice sites crucial to alternative splicing.

## Introduction

Split genes are a defining characteristic of eukaryotic genomes (Plaschka et al., 2019). During pre-mRNA transcription, intervening sequences (introns) are excised, and the flanking sequences (exons) are spliced together. In developmentally simple eukaryotes, such as the experimental model *Saccharomyces cerevisiae*, a relatively small proportion of protein coding genes are split by introns, and the *cis-*element sequences that guide splice site recognition are almost invariant. In developmentally complex eukaryotes, such as humans or Arabidopsis, most protein-coding genes have introns and the *cis*-elements controlling splicing are more degenerate. Such sequence variation facilitates alternative splice site selection that, in turn, permits the regulation of mRNA expression and the production of functionally different protein isoforms (Lee and Rio, 2015; Nilsen and Graveley, 2010). Consistent with this, alternative splicing is the foremost genomic predictor of developmental complexity (Chen et al., 2014).

Pre-mRNA splicing is carried out by the spliceosome (Plaschka et al., 2019; Wilkinson et al., 2020). This large, dynamic molecular machine comprises more than 100 proteins and five uridylate-rich small nuclear RNAs (UsnRNAs). Splicing requires two sequential transesterification reactions (Figure 1). The first reaction, called branching, occurs when the 2’ hydroxyl of the conserved intron branchpoint (BP) adenosine performs a nucleophilic attack on the 5’ splice site (5’SS), resulting in a cleaved 5’ exon and a branched lariat-intron intermediate. The second reaction, exon ligation, occurs via nucleophilic attack of the 3’ hydroxyl of the 5’ exon on the 3’ splice site (3’SS). The recognition of the BP, 5’SS and 3’SS involves ordered base pairing interactions with UsnRNAs and the function of spliceosomal proteins. Typically, the 5’SS is first recognised by U1 snRNP, whilst U2 snRNP recognises the BP sequence of pre-mRNA. The pre-assembled U4/U6.U5 tri-snRNP subsequently joins the spliceosome. The conserved U6 snRNA ACAGA sequence replaces U1 snRNP at the 5’SS and loop 1 of U5 snRNA binds 5’ exon sequences adjacent to the 5’SS (Figure 1). In humans, selection of the 5’SS occurs during this hand-off to U6 and U5 snRNAs, and this step is decoupled from formation of the active site, thus potentially allowing for plasticity of 5’SS selection (Charenton et al., 2019; Fica, 2020). Subsequently, U6 pairs with U2 snRNA and intramolecular U6 snRNA interactions facilitate the positioning of the catalytic metal ions to effect branching and exon ligation (Wilkinson et al., 2020).

**Figure 1.**
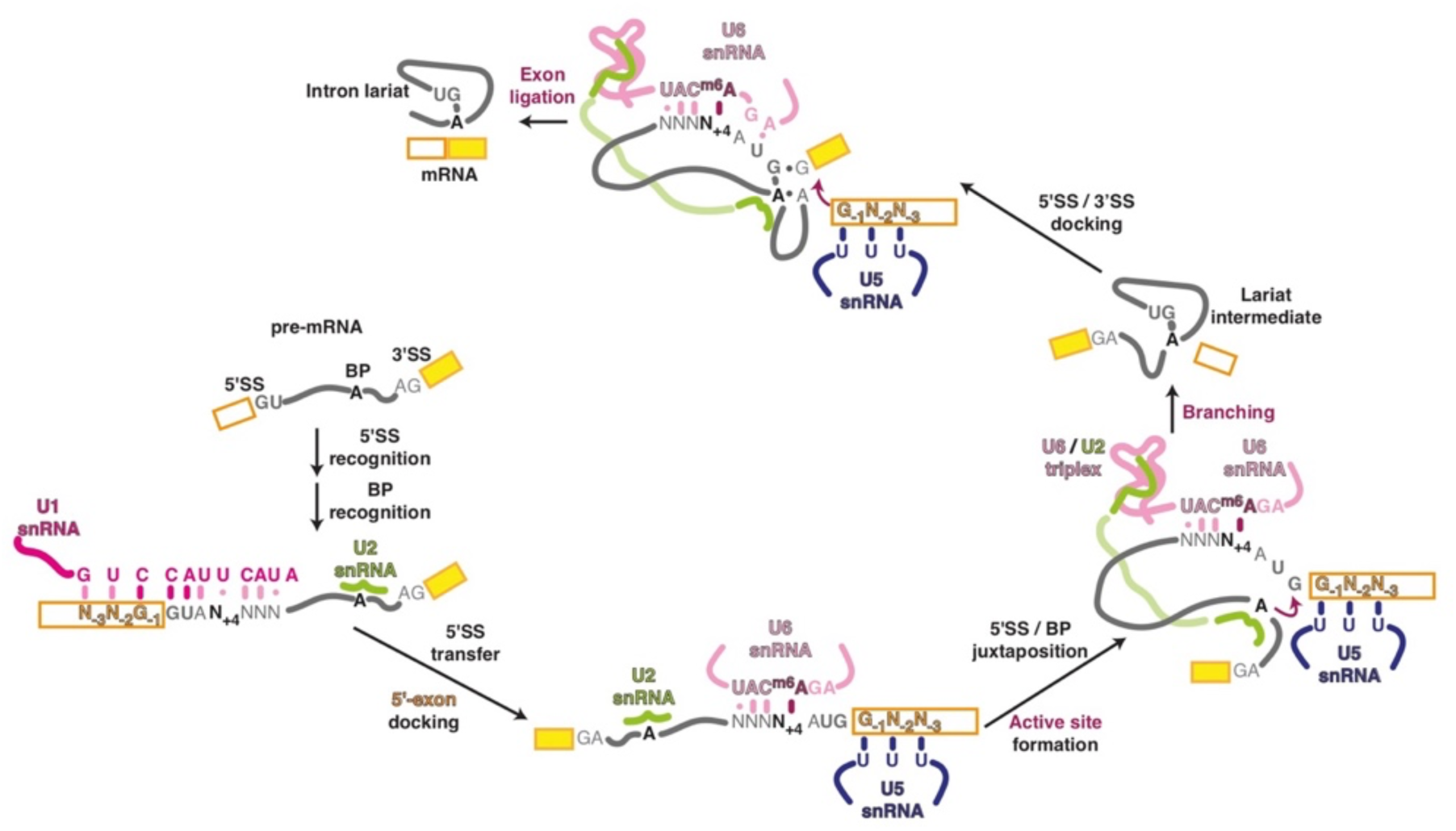
Schematic outline of sequential steps in pre-mRNA splicing. The 5’SS is first recognised by U1 snRNA base-pairing. U2 snRNA recognises the branch point (BP). The 5’SS is transferred to the U6 snRNA ACAGA box (which binds to the intron) and U5 snRNA loop 1 (which docks the upstream exon). The active site then forms through U6/U2 pairing and intramolecular U6 interactions, juxtaposing the selected 5’SS and BP, and enabling nucleophilic attack of the 2’ hydroxyl of the BP A on the 5’SS G+1. The separation of these events in humans provides plasticity for alternative splice site recognition and selection. However, in *S. cerevisiae*, 5’SS transfer and active site formation occur simultaneously. The branching reaction yields the lariat intermediate. Remodelling of the spliceosome for exon ligation involves altered interactions of the U6 snRNA ACAGA box and docking of the 5’SS/3’SS at the active site, promoting nucleophilic attack of the 3’ hydroxyl of the 5’ exon on the 3’SS. The exons are ligated, producing the spliced mRNA, and the intron lariat is released

U6 snRNA is the most highly conserved UsnRNA, reflecting its crucial roles at the active site of the spliceosome (Wilkinson et al., 2020). Different base modifications are found in each of the UsnRNAs, including U6 snRNA, but the role of these modifications is poorly understood (Morais et al., 2021). The essential ACAGA sequence of *S. cerevisiae* U6 snRNA is unmodified. However, in other species including *Schizosaccharomyces pombe* (Gu et al., 1996), the plant *Vicia faba* (Kiss et al., 1987) and human (Shimba et al., 1995), the central adenosine in the corresponding sequence is modified by methylation at the N6 position: AC^m6^**<u>A</u>**GA. In *S. cerevisiae*, U6 snRNA ACAGA recognises the stringently conserved GUAUGU sequence at the 5’ end of introns, with the central adenosine making a Watson-Crick base pair with the almost invariant U at the +4 position of the 5’SS (5’SS U_+4_) (Neuvéglise et al., 2011; Wan et al., 2019). Conversely, in species with m^6^A modified U6 snRNA the 5’SS is degenerate and the identity of the base at the +4 position varies, but is usually enriched for A. Therefore, the pairing between the 5’SS and the U6 snRNA AC**<u>A</u>**GA box is unlikely to be driven primarily by canonical Watson-Crick base pairs in these species. The precise function of the U6 snRNA m^6^A modification is unknown.

In humans, the U6 snRNA AC**<u>A</u>**GA sequence is methylated by the conserved methyltransferase METTL16 (Aoyama et al., 2020; Pendleton et al., 2017; Warda et al., 2017). This modification depends upon specific sequence and distinct structural features of U6 snRNA that are recognised by METTL16. Hairpin sequences that mimic these features of U6 snRNA are found within the 3’UTR of S-adenosylmethionine (SAM) synthetase *MAT2A* mRNA and are also methylated by METTL16 (Pendleton et al., 2017; Shima et al., 2017). SAM synthetase is the enzyme responsible for production of the methyl donor SAM, which is required for methylation reactions in the cell. Binding of METTL16 to *MAT2A* mRNA influences splicing and/or stability, regulating MAT2A expression and SAM levels (Pendleton et al., 2017; Shima et al., 2017). Mammalian METTL16 knock-outs are embryo lethal (Mendel et al., 2018). In *Caenorhabditis elegans*, the METTL16 orthologue, METT-10, methylates U6 snRNA and also influences SAM levels by targeting SAM synthetase (*sams*) genes (Mendel et al., 2021). METT-10 methylates the 3’SS of *sams-3/4* intron 2, although in a different sequence and structural context to U6 snRNA (Mendel et al., 2021). In *S. pombe*, the METTL16 orthologue, MTL16, appears to target only U6 snRNA (Ishigami et al., 2021). *S. pombe* mutants defective in MTL16 function show less efficient splicing of some introns, resulting in increased levels of intron retention. The *S. pombe* introns sensitive to loss of MTL16 function are distinguished by having adenosine at the +4 position of the intron (5’SS A_+4_). However, the effect of 5’SS A_+4_ is partially suppressed in introns that have stronger predicted base pairing between U5 snRNA loop 1 and the 5’ exon (Ishigami et al., 2021).

Alternative splicing plays crucial roles in gene regulation, affecting diverse biological processes including the control of flowering in response to ambient temperature (Airoldi et al., 2015; Capovilla et al., 2017). Plants control the time at which they flower by integrating responses to environmental cues, such as temperature and daylength, with an endogenous program of development (Andrés and Coupland, 2012). Slight changes in ambient temperature can profoundly influence flowering time. Indeed, documented changes in the flowering times of many plant species have provided some of the best biological evidence of recent climate change (Fitter and Fitter, 2002). At cooler ambient temperatures, pre-mRNAs encoding repressors of Arabidopsis flowering, FLM (FLOWERING LOCUS M) and MAF2 (MADS AFFECTING FLOWERING 2), are efficiently spliced (Airoldi et al., 2015; Balasubramanian et al., 2006; Lee et al., 2013; Lutz et al., 2015; Posé et al., 2013; Scortecci et al., 2001). As a result, FLM and MAF2 proteins are produced at cooler temperatures and form higher order protein complexes with other MADS box factors, such as MAF3 (*MADS AFFECTING FLOWERING 3*), SVP (*SHORT VEGETATIVE PHASE*) and FLC (*FLOWERING LOCUS C*), to directly repress the expression of the genes *FT (FLOWERING LOCUS T)* and *SOC1 (SUPPRESSOR OF OVEREXPRESSION OF CO 1)*, which promote flowering (Gu et al., 2013). At elevated ambient temperatures, the splicing of *MAF2* and *FLM* introns is adjusted, in different ways, effecting increased levels of non-productive transcripts (Airoldi et al., 2015; Balasubramanian et al., 2006; Lee et al., 2013; Lutz et al., 2015; Posé et al., 2013; Scortecci et al., 2001). Consequently, the abundance of active MAF2 and FLM proteins is compromised at elevated temperatures: repression of genes promoting flowering is relieved, and flowering is enabled (Airoldi et al., 2015; Capovilla et al., 2017; Posé et al., 2013; Rosloski et al., 2013). In the case of *MAF2*, this temperature-responsive splicing is characterised by a progressive increase in retention of intron 3 as ambient temperature increases (Airoldi et al., 2015). The thermosensory mechanism that underpins the temperature-responsive alternative splicing of *MAF2* and *FLM* is unknown.

We conducted a mutant screen for factors required for the efficient splicing of *MAF2* pre-mRNA intron 3 at cooler temperatures. This screen identified an early flowering mutant disrupting the gene encoding the Arabidopsis METTL16 orthologue, FIONA1 (FIO1) (Kim et al., 2008). We show that FIO1 is required for m^6^A modification of U6 snRNA, consistent with recent reports confirming this conserved function (Wang et al., 2022; Xu et al., 2022). We reveal the impact of FIO1 on *MAF2* splicing and global patterns of gene expression. We detect widespread changes in pre-mRNA splicing in *fio1* mutants that can be explained by the identity of the base at the 5’SS +4 position of affected introns, and by cooperative roles for U5 and U6 snRNA in 5’SS selection. We also reveal that U6 snRNA interactions at 5’SSs affect 3’SS choice. Analysis of other annotated genomes reveal the existence of two major classes of 5’SS. Our findings suggest cooperative and compensatory roles of U5 and U6 snRNA may contribute to 5’SS selection in a range of eukaryotes.

## Results

### Identification of *FIO1* in a screen for early flowering mutants with increased retention of *MAF2* intron 3

To identify factors responsible for promoting efficient splicing of *MAF2* intron 3 at low ambient temperature, we carried out a mutant screen using ethyl methanesulfonate (EMS) as a mutagen. We screened for Arabidopsis mutants that flowered early at 16°C. The earliest flowering 100 individuals were then re-screened for enhanced *MAF2* intron 3 retention at 16°C, using RT-PCR. Two independent mutants (EMS-129 and EMS-213) showed early flowering and increased levels of *MAF2* intron 3 retention at 16°C.

We used bulked segregant analysis and the software tool artMap (Javorka et al., 2019) to identify the causative mutation in EMS-129 as a G to A transition at position 9,041,454 of chromosome 2, which disrupts the 5’SS of AT2G21070 intron 2. A mutation in AT2G21070 has previously been described and named *fiona1-1* (Kim et al., 2008). Crosses between EMS 129 and either *fio1-1*, or a Transfer-DNA insertion line disrupting AT2G21070 (SALK_084201); *fio1-3*) confirmed allelism. We therefore refer to EMS-129 as *fio1-4*. We summarize the molecular basis of the three alleles in Figure 2A. The disruption of the 5’SS of intron 2 in *fio1-4* results in activation of an upstream cryptic 5’SS within exon 2 and the excision of 69 nt from the coding region of the mRNA (Figure 2A). We found that the *fio1-1* and *fio1-3* alleles also show increased levels of *MAF2* intron 3 retention at low temperature (Figure 2B) and like *maf2* mutants, flower early (Figure 2C-D). *fio1* alleles display additional visible phenotypes, including reduced apical dominance and stature (Figure 2D). *FIO1* encodes the Arabidopsis orthologue of the human methyltransferase, METTL16 (Kim et al., 2008; Pendleton et al., 2017).

**Figure 2.**
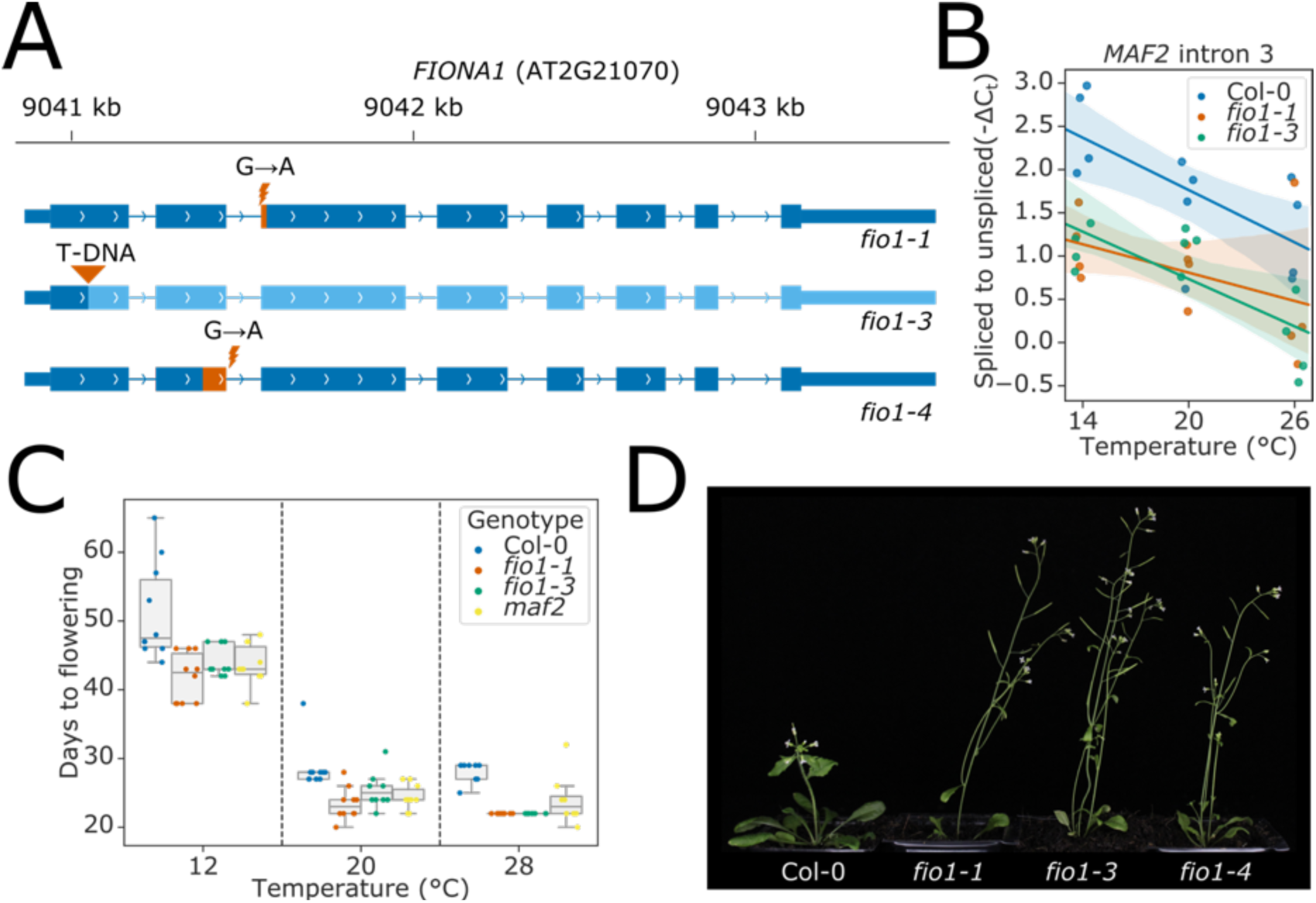
Loss of *FIO1* causes early flowering and reduced splicing of pre-mRNA encoding the floral repressor MAF2: **(A)** Gene track showing the three *fio1* mutant alleles used in this study. *fio1-1* is an EMS mutant with a G→A transition at the -1 position of the 3’SS in intron 2 of *FIO1* (Kim et al., 2008). This causes activation of a cryptic 3’SS 15 nt downstream, and the loss of 5 aa of sequence from the *FIO1* open reading frame (shown in orange). *fio1-3* is a T-DNA insertion mutant (SALK_084201) in the first exon of the *FIO1,* disrupting the gene (region downstream of insertion shown in light blue). *fio1-4* is an EMS mutant with a G→A transition at the +1 position of the 5’SS in intron 2 of *FIO1*. This causes activation of a cryptic 5’SS 69 nt upstream, and the loss of 23 aa of sequence from the *FIO1* open reading frame (shown in orange). **(B)** Regression scatterplot showing the change in spliced to retained ratio of *MAF2* intron 3 in *fio1-1* and *fio1-3* at a range of temperatures. Shaded regions show bootstrapped 95% confidence intervals for regression lines. **(C)** Boxplot showing the change in flowering time observed in the *fio1-1, fio1-3* and *maf2* mutants at a range of temperatures. **(D)** Photographs showing the early flowering phenotypes of *fio1-1*, *fio1-3* and *fio1-4* mutants.

### FIO1-dependent methylation of poly(A)+ RNA is rare or absent

We first asked which RNAs were methylated by FIO1. We have previously used nanopore direct RNA sequencing (DRS) to map m^6^A sites dependent on the Arabidopsis METTL3-like writer complex component VIRILIZER (VIR), revealing m^6^A enriched in a DR^m6^**A**CH consensus in the 3’ terminal exon of protein coding mRNAs (Parker et al., 2020). We used a similar nanopore DRS experiment to map FIO1-dependent m^6^A, sequencing poly(A)+ RNA purified from wild-type (Col-0) and *fio1-1*. As a positive control, we sequenced poly(A)+ RNA purified from the *fip37-4* mutant, which is defective in the Arabidopsis orthologue of the METTL3 m^6^A writer complex component WTAP (Zhong et al., 2008). We sequenced four biological replicates of each genotype, resulting in a total of 22.7 million mapped reads. The corresponding sequencing statistics are detailed in Supplementary file 1.

We used the software tool Yanocomp to map differences in mRNA modifications in Col-0, *fio1-1* and *fip37-4* (Parker et al., 2021a). Applying this approach, we identified 37,861 positions that had significantly different modification rates in a three-way comparison between Col-0, *fip37-4* and *fio1-1* (Figure 3A). Of these, 97.9% had significant changes in modification rate in a pairwise-comparison between *fip37-4* and Col-0. In contrast, only 7.6% had altered modification rates in *fio1-1*. Of these, 85.8% also had altered modification rates in *fip37-4* (Figure 3A), with larger effect sizes (Figure 3 – figure supplement 1A), indicating that they are METTL3-like writer complex-dependent m^6^A sites whose modification rate is indirectly affected by loss of FIO1. The FIP37-dependent modification sites, including those with small effect size changes in *fio1-1,* were found in a DRACH consensus in 3’UTR regions (Figure 3 – figure supplement 1B-C), consistent with established features of m^6^A modifications deposited by the Arabidopsis METTL3-like writer complex (Parker et al., 2020).

**Figure 3.**
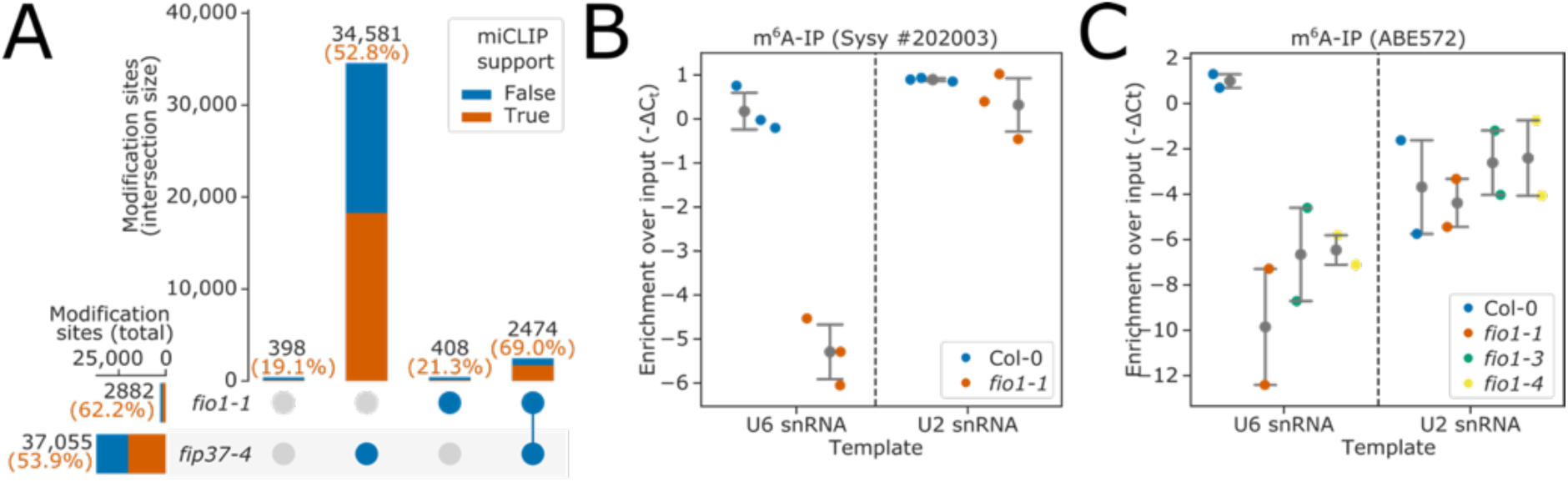
FIO1-dependent m^6^A modification of poly(A)+ mRNA is rare or absent but FIO1 is required for U6 snRNA methylation. **(A)** Upset plot of m^6^A modification sites detected by Yanocomp (Parker et al., 2021a). All sites shown have significant differences in modification level in a three-way comparison between *fip37-4, fio1-1,* and Col-0. Bars show size of intersections between sites which are significant in each two-way comparison. Total intersection sizes displayed in black above each bar. Orange and blue bar fractions show the number of sites within each set intersection that have or do not have an miCLIP peak (Parker et al., 2020) within 5 nt, respectively. Percentage of intersections with miCLIP support are displayed in orange above each bar. A small number of sites are significant in the three-way comparison, but in neither two-way comparison (far left bar). **(B-C)** Stripplots with mean and 95% confidence intervals, showing the enrichment of U6 and U2 snRNAs over input using **(B)** Synaptic systems #202 003 anti-m^6^A antibody and RNA purified from Col-0 and *fio1-1*, and **(C)** Millipore ABE572 anti-m^6^A antibody and RNA purified from Col-0, *fio1-1*, *fio1-3* and *fio1-4.* Y axes show — ΔC_t_ (m^6^A-IP — input) corrected for input dilution factor. Striplots show mean values for three or four independent RT-qPCR amplifications on each biological replicate immunopurification experiment.

We used the antibody-based technique miCLIP to perform orthogonal validation of predicted modification sites (Parker et al., 2020): *FIP37-*dependent m^6^A sites were well supported, with 52.8% less than 5 nt from an miCLIP peak (Figure 3A). In contrast, of the 408 sites discovered only in *fio1-1*, just 21.3% (87) were less than 5 nt from an miCLIP peak, indicating that the majority are likely to be false positives. The 87 positions with miCLIP support were found in a DRACH consensus in 3’UTRs, suggesting that they are false negative *FIP37-*dependent m^6^A sites rather than genuine FIO1-dependent sites (Figure 3 – figure supplement 1D-E).

We did not find *FIO1-*dependent methylation changes in transcripts of the four Arabidopsis homologues of SAM-synthetase (Figure 3 – figure supplement 2A-D). Consequently, the METTL16-dependent regulation of SAM homeostasis identified in metazoans (Mendel et al., 2021; Pendleton et al., 2017; Shima et al., 2017) appears not to be conserved in Arabidopsis. Consistent with previous evidence that m^6^A in 3’UTRs affects cleavage and polyadenylation (Parker et al., 2020), we identified 1104 genes with altered poly(A) site choice in *fip37-4,* compared to only 49 in *fio1-1* (Figure 3 – figure supplement 1F). Overall, these findings indicate that FIO1-dependent m^6^A sites in poly(A)+ mRNA are rare or absent.

### m^6^A modification of U6 snRNA depends upon FIO1

Since FIO1 orthologues methylate U6 snRNA, we asked if FIO1 was required to modify Arabidopsis U6 snRNA. We performed two experiments using two independent anti-m^6^A antibodies to immunopurify methylated RNAs from Col-0 and *fio1* alleles. For control, we used U2 snRNA, which is m^6^A methylated in humans, but not by METTL16 (Chen et al., 2020). Using RT-qPCR, we could detect equivalent levels of U6 snRNA in the input RNA purified from different genotypes, suggesting that the abundance or stability of U6 snRNA is unaffected by loss of FIO1 function (Figure 3 – figure supplement 4A-B). We could detect enrichment of U6 and U2 snRNAs in RNA immunopurified with anti-m^6^A antibodies from Col-0 (Figure 3B-C). The enrichment of U2 snRNA in these experiments was unaffected by loss of FIO1 function. In contrast, we identified significant depletion of U6 snRNA in the anti-m^6^A immunopurified RNA from *fio1-1, fio1-3* and *fio1-4* alleles (Figure 3B-C). These data are consistent with recent reports (Wang et al., 2022; Xu et al., 2022) indicating that Arabidopsis U6 snRNA is m^6^A-modified at the conserved position of the AC^m6^**<u>A</u>**GA box and that this modification requires active FIO1.

### Widespread disruption of pre-mRNA splicing in *fio1* mutants

Having found that FIO1 is required for U6 snRNA m^6^A modification, we examined changes in gene expression and pre-mRNA splicing in *fio1* mutants. Nanopore DRS is insightful for mapping the complexity of RNA processing and modification, but throughput currently limits the statistical power to detect changes. Splicing analysis using nanopore DRS is also currently limited by basecalling error rate, which complicates the alignment of exon/intron boundaries (Parker et al., 2021b). Consequently, we used Illumina RNA-Seq of poly(A)+ RNA to analyse gene expression and splicing. Since FIO1 affects the splicing of *MAF2*, which is sensitive to changes in temperature, we included plants grown at 20°C and shifted to either 4°C, 12°C, or 28°C for 4h prior to harvesting and poly(A)+ RNA purification. We sequenced six biological replicates each of wild-type Col-0 and *fio1-3*, generating a minimum of 46 million paired end reads, 150 bp in length, per replicate. On average, 97.9% of read pairs were mappable per replicate, resulting in a total of 3.6 billion mapped read pairs. A summary of the sequencing read statistics is given in Supplementary file 1. The RNA-Seq data clearly reveals disruption of *FIO1* gene expression in *fio1-3* (Figure 4 – figure supplement 1A).

In order to detect cryptic splice sites that might be activated in *fio1* mutants, but not annotated in available Arabidopsis reference transcriptomes, we built a bespoke reference transcriptome derived from the Col-0/*fio1-3* Illumina RNA-Seq and Col-0/*fio1-1* nanopore DRS data using the software tool Stringtie2 (Kovaka et al., 2019). We quantified the expression of these transcripts using Salmon (Patro et al., 2017), then used SUPPA2 to calculate event-level percentage changes in splicing, which also classifies different types of splicing event (Trincado et al., 2018). These percent spliced indices (PSI) were used to fit linear models for each splicing event to identify changes in splicing dependent on temperature, genotype, and temperature × genotype interactions. Analysis of the Col-0 temperature-shift data served as a control. Consistent with previous studies (Calixto et al., 2018), we found that intron retention is the predominant class of alternative splicing event detected when Arabidopsis is subjected to different temperatures (Figure 4A). In contrast, when we repeated this analysis to classify alternative splicing events that differ between *fio1-3* and Col-0, we found that a larger proportion of alternative splicing events were classified as alternative 5’SS usage – 34.4%, compared to only 18.6% of temperature-dependent events (Figure 4B). In addition, we detected changes in the PSI of retained introns, exon skipping and alternative 3’SS selection (Figure 4B). There was a significant overlap between the alternative splicing events that were sensitive to loss of *FIO1,* and those that were sensitive to temperature (hypergeometric-test p < 1×10^—16^). However, in 64.2% of *fio1-*sensitive events, loss of *FIO1* did not alter splicing responses to temperature (Figure 4B). Of the remaining 2505 splicing events that did have altered temperature sensitivity in the absence of FIO1, 38.4% were alternative 5’SSs, and of these, 69.9% had greater sensitivity to loss of *FIO1* at 28°C than at 4°C. This suggests that FIO1 enables accurate 5’SS selection at elevated temperature (Figure 4 – figure supplement 1A).

**Figure 4.**
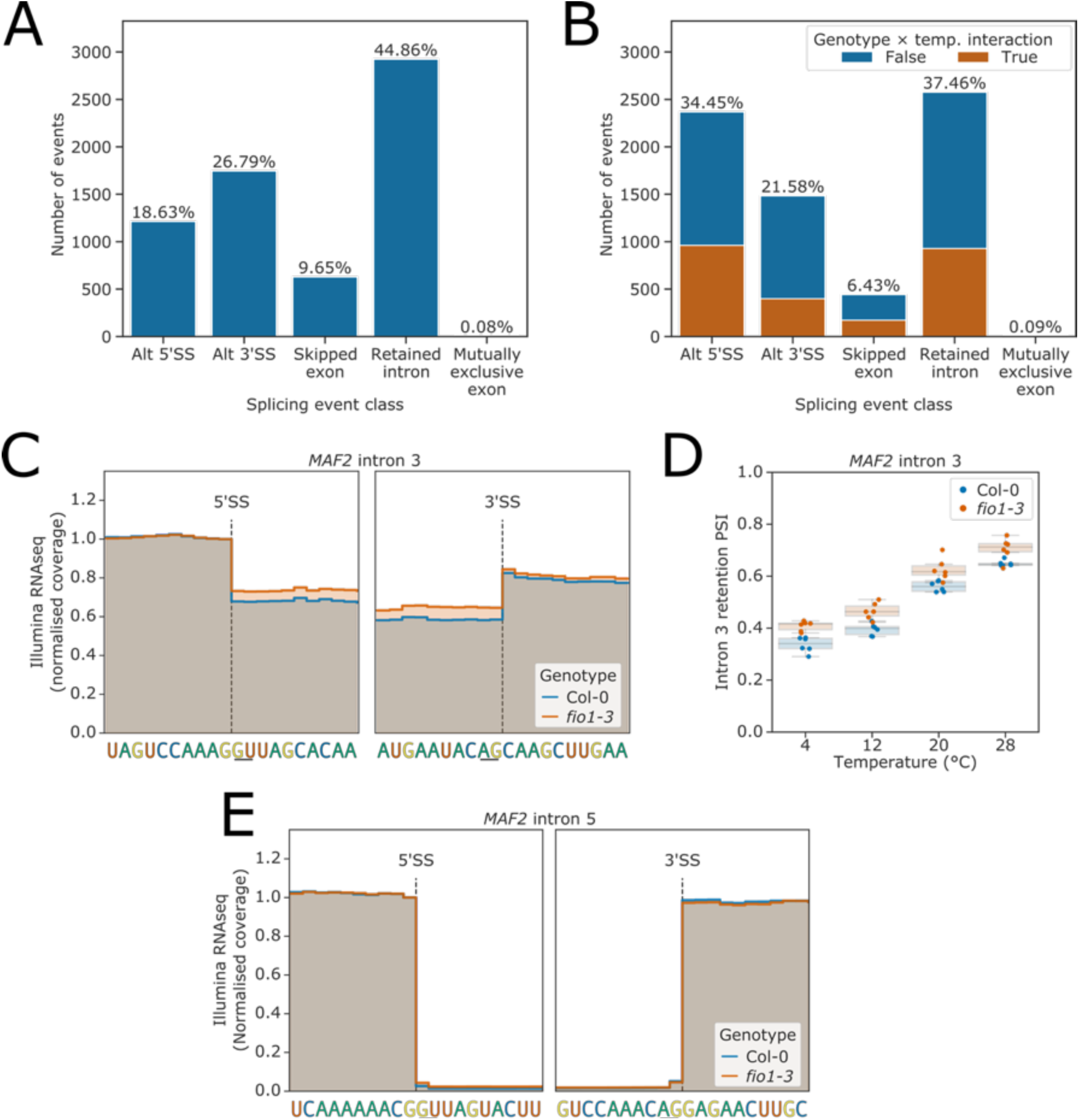
Splicing events sensitive to temperature and loss of *FIO1.* **(A-B)** Barplots showing the proportion of splicing events of each class, as labelled by SUPPA, which have significantly different usage (FDR < 0.05) at either **(A)** varying temperatures or **(B)** in *fio1-3*. In **(B)**, events for which the response to temperature changes in *fio1-3* are shown in orange. **(C)** Gene track showing the change in retention of *MAF2* intron 3 in *fio1-3*, at 20°C. Expression is normalised by the coverage at the —1 position of the 5’SS. **(D)** Boxplot showing the change in retention of *MAF2* intron 3 at varying temperatures and in *fio1-3*. **(E)** Gene track showing the absence of a change in splicing of *MAF2* intron 5 in *fio1-3*, at 20°C. Expression is normalised by the coverage at the —1 position of the 5’SS.

Analysis of the RNA-Seq data confirmed the increased retention of *MAF2* intron 3 in *fio1-3,* consistent with the results of our mutant screen (Figure 2B & 4C). Loss of *FIO1* decreased the splicing efficiency of *MAF2* intron 3 by approximately equivalent amounts at all temperatures (Figure 4D), implying that FIO1 is not required to generate *MAF2* intron 3 temperature sensitivity, but calibrates the range over which it occurs. Loss of *FIO1* had no effect on the splicing of *MAF2* intron 5, indicating that this effect is specific to particular *MAF2* introns (Figure 4E). We could also detect changes in gene expression consistent with the early flowering phenotype of *fio1* mutant alleles (Figure 4 – figure supplement 1C-I). For example, the mRNA expression levels of *FT* and *SOC1* were increased in *fio1-3* (Figure 4 – figure supplement 1C-D). In contrast, the expression levels of the floral repressors *FLM*, *MAF2, MAF3, MAF4, MAF5* and *FLC* were reduced (Figure 4 – figure supplement 1E-J). In the case of *FLC*, both sense and antisense transcript levels were reduced in *fio1-3* (Figure 4 – figure supplement 1J-K), but there were no detectable changes in splicing patterns, nor in the alternative polyadenylation of the antisense transcripts (Figure 4 – figure supplement 1L). In these experimental conditions, loss of FIO1 did not affect RNA-level expression or splicing of either *CO* or *SVP* (Figure 4 – figure supplement 1M-N). In contrast, *fio1-3* mutants exhibited defects in the splicing of pre-mRNA encoding MAF2, MAF3, FLM, and other genes that influence flowering time, as detailed further below.

Overall, we conclude that the predominant molecular phenotype resulting from loss of FIO1 function is a major disruption to patterns of splicing, in a manner that is mostly independent of temperature. To understand the basis of these splicing differences, we analysed each of these splicing classes separately.

### Alternative selection of 5’SSs in *fio1-3* can be explained by U6 and U5 snRNA target sequences

We identified 2369 changes in 5’SS choice caused by loss of FIO1 function. We asked if differences in *cis*-element features could account for FIO1-dependent 5’SS selection. Comparing 5’SSs that exhibited reduced selection in *fio1-3* with their corresponding alternative 5’SSs, which were more frequently selected in *fio1-3,* we found a significant difference in base composition at the —2 to +5 positions (G-test p < 1×10^—16^). Specifically, we identified a //GURAG motif (R = A or G) at 5’SSs sensitive to the loss of FIO1 function (Figure 5A). This motif is consistent with recognition by the U6 snRNA ACAGA box (Kandels-Lewis and Séraphin, 1993; Lesser and Guthrie, 1993; Sawa and Abelson, 1992). In contrast, the alternative 5’SSs selected more frequently in *fio1-3* were characterised by an AG//GU motif (Figure 5B). This motif is consistent with U5 snRNA loop 1 recognition of the upstream exon sequences (Galej et al., 2016; Newman and Norman, 1992). A heat map of the sequence composition of *fio1-3*-sensitive 5’SSs reveals that they are more likely to be characterised by strong base-pairing interactions with the U6 snRNA ACAGA box, but weak interactions with U5 snRNA loop 1 (Figure 5A). In contrast, a heat map of the composition of 5’SS sequences that are preferentially used in *fio1-3,* shows strong interactions with U5 snRNA loop 1, but weak interactions with the U6 snRNA ACAGA box (Figure 5B).

**Figure 5.**
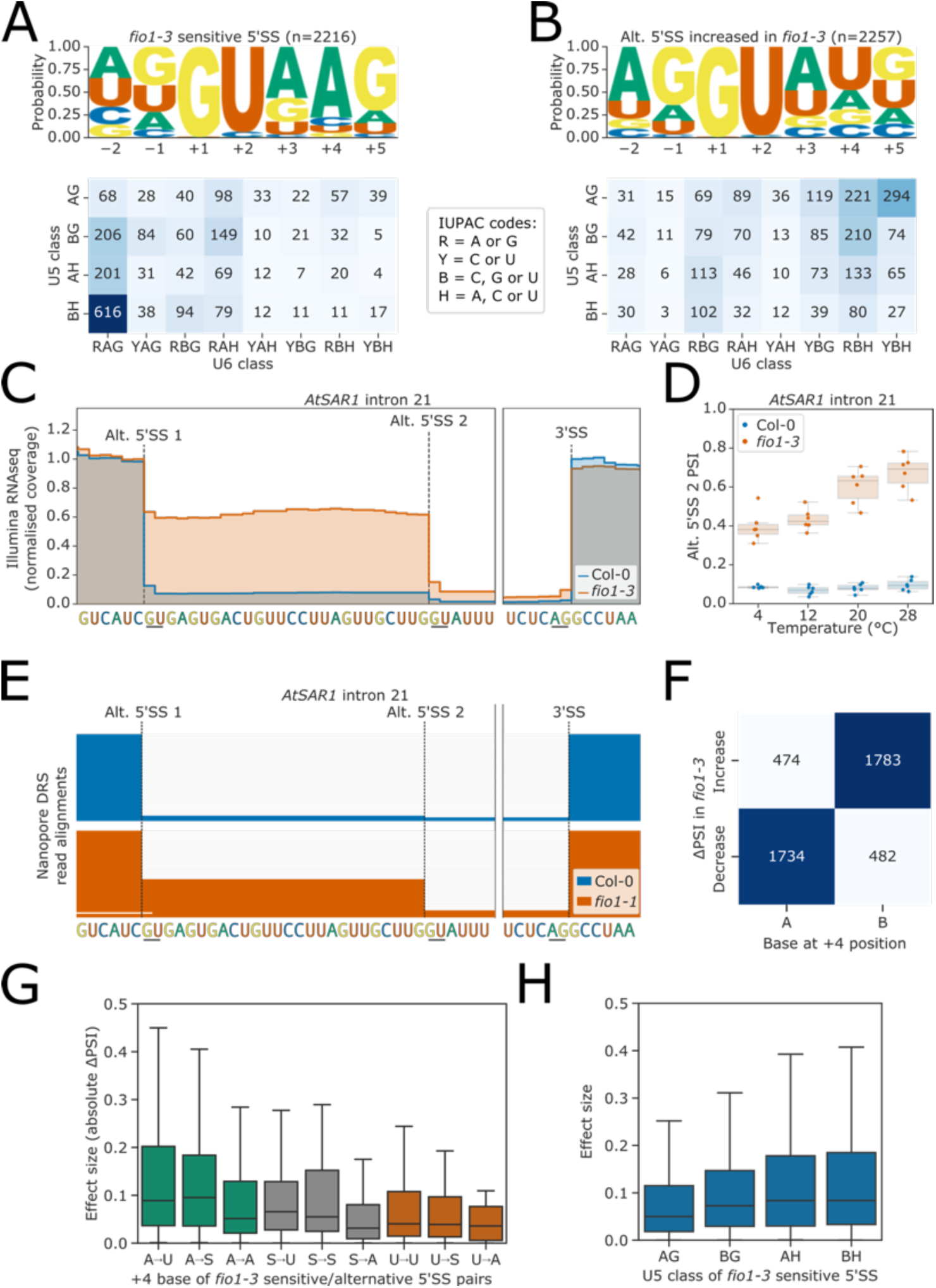
Effect of *fio1-3* on alternative 5’ splice sites. **(A-B)** Sequence logos and heatmap showing the distribution of U5 snRNA and U6 snRNA interacting sequence classes for 5’SSs which **(A)** are sensitive to loss of *FIO1* function or **(B)** have increased usage in *fio1-3.* Motifs are shown for the —2 to +5 positions of the 5’SS. U5 classes are based upon the distance of the —2 to —1 positions of the 5’SS from the consensus motif AG. U6 classes are based upon the distance of the +3 to +5 positions of the 5’SS from the consensus motif RAG. **(C)** Gene track showing alternative 5’SS usage at *AtSAR1* intron 21 in *fio1-3*, at 20°C. Expression is normalised by the coverage at the —1 position of the 5’SS. **(D)** Boxplot showing the change in usage of the cryptic alternative 5’SS (Alt 5’SS 2) in *AtSAR1* intron 21 at varying temperatures, in Col-0 and *fio1-3*. **(E)** Gene track showing alternative 5’SS usage at *AtSAR1* intron 21 in *fio1-1*, identified using nanopore DRS read alignments. Alignments have been subsampled to a maximum of 50 per condition. **(F)** Contingency table showing the relationship between the nucleotide at the +4 position, and the direction of change in 5’SS usage in *fio1-3,* for pairs of alternative 5’SSs with significantly altered usage in *fio1-3.* **(G)** Boxplot showing effect sizes of pairs of alternative 5’SSs with significantly altered usage in *fio1-3*, separated by +4 position bases (A→U indicates that 5’SS with reduced usage has A_+4_, 5’SS with increased usage has U_+4_). **(H)** Boxplot showing effect sizes of pairs of alternative 5’SSs with significantly altered usage in *fio1-3*, separated by U5 classification

The widespread changes in 5’SS selection are evident at individual loci. For example, at *AtSAR1* (AT1G33410), which encodes a nucleoporin (Parry et al., 2006), loss of FIO1 function at 20°C results in a 52.2% reduction in the use of the normally near-constitutive 5’SS in intron 21, with sequence UC//GUGAG (Figure 5C, D). There is a reciprocal increase in the use of a cryptic 5’SS 26 nt downstream, with sequence UG//GUAUU. At elevated temperatures, use of this cryptic 5’SS becomes increasingly preferred (Figure 5D). The change in usage of this 5’SS is also visible in the *fio1-1* allele mapped using orthogonal nanopore DRS (Figure 5E). At *MAF2*, in addition to changes in intron 3 retention, we also detected decreased use of the canonical 5’SS at intron 2 (UA//GUAAG), and increased use of a cryptic 5’SS in intron 2 (UC//GUACU; Figure 5 – figure supplement 1A-B). At another floral repressor gene, *FLM* (AT1G77080), which has mutually exclusive splicing of exons 2 and 3, we found that 5’SSs at the 3’ ends of both exon 2 (UA//GUAAG) and exon 3 (UC//GUAAC) had reduced usage in *fio1-3*, with a concomitant increase in the use of a cryptic 5’SS within exon 3 (AU//GUUUU; Figure 5 – figure supplement 1C-D). The overall effect of these (and other) splicing changes is that the fraction of productive *AtSAR1*, *MAF2* and *FLM* transcripts is reduced in *fio1-3* (Figure 5 – figure supplement 1E-G). Finally, at the gene encoding the METTL3-like writer complex component MTB (the orthologue of human METTL14), loss of FIO1 function resulted in a 40.4% increase in the use of a cryptic 5’SS in intron 4 that introduces a premature termination codon into the *MTB* open reading frame (Figure 5 – figure supplement 1H-I). Consequently, loss of *FIO1* may have indirect effects on m^6^A deposition by the METTL3-like writer complex.

Remarkably, the presence or absence of an A at the +4 position (A_+4_) correctly separates 78.6% of the 5’SSs exhibiting decreased and increased usage in *fio1-3* (Figure 5F). In humans, where pairing of the 5’SS to U6 snRNA occurs before activation of the spliceosome, the 5’SS A_+4_ faces the m^6^A of the U6 snRNA AC^m6^**<u>A</u>**GA box in the B complex before docking of the 5’SS in the active site (Bertram et al., 2017). Of the 5’SSs with increased usage in *fio1-3*, 48.2% have U_+4_, which could make a Watson–Crick base pair with the corresponding unmethylated residue of U6 snRNA. In total, 61.8% of 5’SS changes in *fio1-3* are associated with a switch from A_+4_ to B_+4_ (A→B_+4_, B = C, G or U), of which 58.8% are A→U_+4_ (Figure 5 – figure supplement 2A). In comparison, only 3.8% of alternative 5’SS pairs are reciprocal B→A_+4_ switches, indicating that this shift is strongly directional. A further 8.0% of alternative 5’SS pairs are S→U_+4_, suggesting that a Watson–Crick A-U base-pair is favoured when U6 snRNA is not m^6^A modified. Surprisingly, 22.3% of alternative 5’SS pairs have the same base at the +4 position (Figure 5 – figure supplement 2A). However, of these 5’SS pairs, the majority are associated with a G→H_+5_ (H = A, C or U) and/or H→G_–1_ switch that weakens interactions with the U6 snRNA ACAGA box and strengthens U5 snRNA loop 1 interactions (Figure 5 – figure supplement 2B).

We next examined how the effect size (absolute ΔPSI) of splicing changes correlated with the base at the +4 position of the 5’SS. The largest effect sizes were associated with A→U_+4_ (Figure 5G). In contrast, alternative 5’SS pairs where there was an A_+4_→A shift had smaller effect sizes. In addition, we found that *fio1-3*-sensitive 5’SSs with 5’SS G_+5_ had larger effect sizes, but that G_+5_ at the alternative 5’SS had less effect, suggesting that G_+5_ is only deleterious in *fio1-3* when in combination with A_+4_ (Figure 5 – figure supplement 2C). Finally, we found that *fio1-3-*sensitive 5’SSs with AG//GU motifs had smaller effect sizes, indicating that favourable interactions with U5 snRNA loop 1 are able to suppress the effect of unfavourable U6 snRNA interactions in *fio1-3* (Figure 5 – figure supplement 2C).

We investigated if there was directionality to shifts in splice site choice in *fio1-3*. Alternative 5’SSs were almost equally likely to be selected either upstream or downstream (Figure 5 – figure supplement 2D). We detected many examples of 5’SS shifts of exactly - 4 nt, +4 nt, and +5 nt. Notably, in +5 nt switches, 5’SSs with the strong U6 snRNA ACAGA recognition sequence //GURAGGU can become strong U5 snRNA loop 1 5’SSs with the sequence GURAG//GU (Figure 5 – figure supplement 2E), suggesting that overlaps in the registers of consensus sequences for U5 and U6 snRNA interactions could facilitate alternative splicing.

We analysed the temperature-dependent shifts in splice site choice in the Col-0 RNA-Seq data as a control. In contrast to the clear difference in sequence composition of 5’SSs with altered use in *fio1-3*, there was no significant difference between the positional base frequencies for 5’SSs that were used preferentially at lower and elevated temperatures in Col-0 (G-test p = 0.28; Figure 5 – figure supplement 2F-G). We conclude that the features of 5’SSs sensitive to loss of FIO1 are specific, and not a generic feature of disrupted splicing patterns.

In summary, these analyses demonstrate widespread change in 5’SS selection in *fio1-3* mutants. The 5’SSs with a strong match to the U6 snRNA ACAGA box and/or an A_+4_ are most sensitive to loss of FIO1. This sensitivity can be suppressed by a strong match in the 5’ exon to U5 snRNA loop 1. Alternative 5’SSs that are used more in *fio1-3* have U_+4_, as well as 5’ exon sequence features that favour recognition by U5 snRNA loop 1.

### FIO1-dependent changes in intron retention and exon skipping can be explained by U6 and U5 snRNA target sequences

In addition to altered 5’SS selection, we detected many instances of intron retention and exon skipping in *fio1-3*. We therefore asked if these FIO1-sensitive splicing phenotypes were associated with specific sequence motifs.

We identified 2576 introns with altered levels of retention in *fio1-3*, of which 55.8% had increased retention and 44.2% had decreased retention (or more splicing). Analysis of 5’SS sequences at introns with increased retention indicates that 76.9% had A_+4_, as well as having weaker interaction potential with U5 snRNA loop 1 (Figure 6A). For example, at *WNK1* (AT3G04910), we detected increased retention of intron 5 (CG//GUGAG) in *fio1-3* (Figure 6B-C) and *fio1-1* (Figure 6D) using Illumina RNA-seq and nanopore DRS analysis, respectively. This indicates that these introns are normally recognised by strong U6 snRNA interactions and are less efficiently spliced in the absence of *FIO1*-dependent U6 snRNA m^6^A modification.

**Figure 6.**
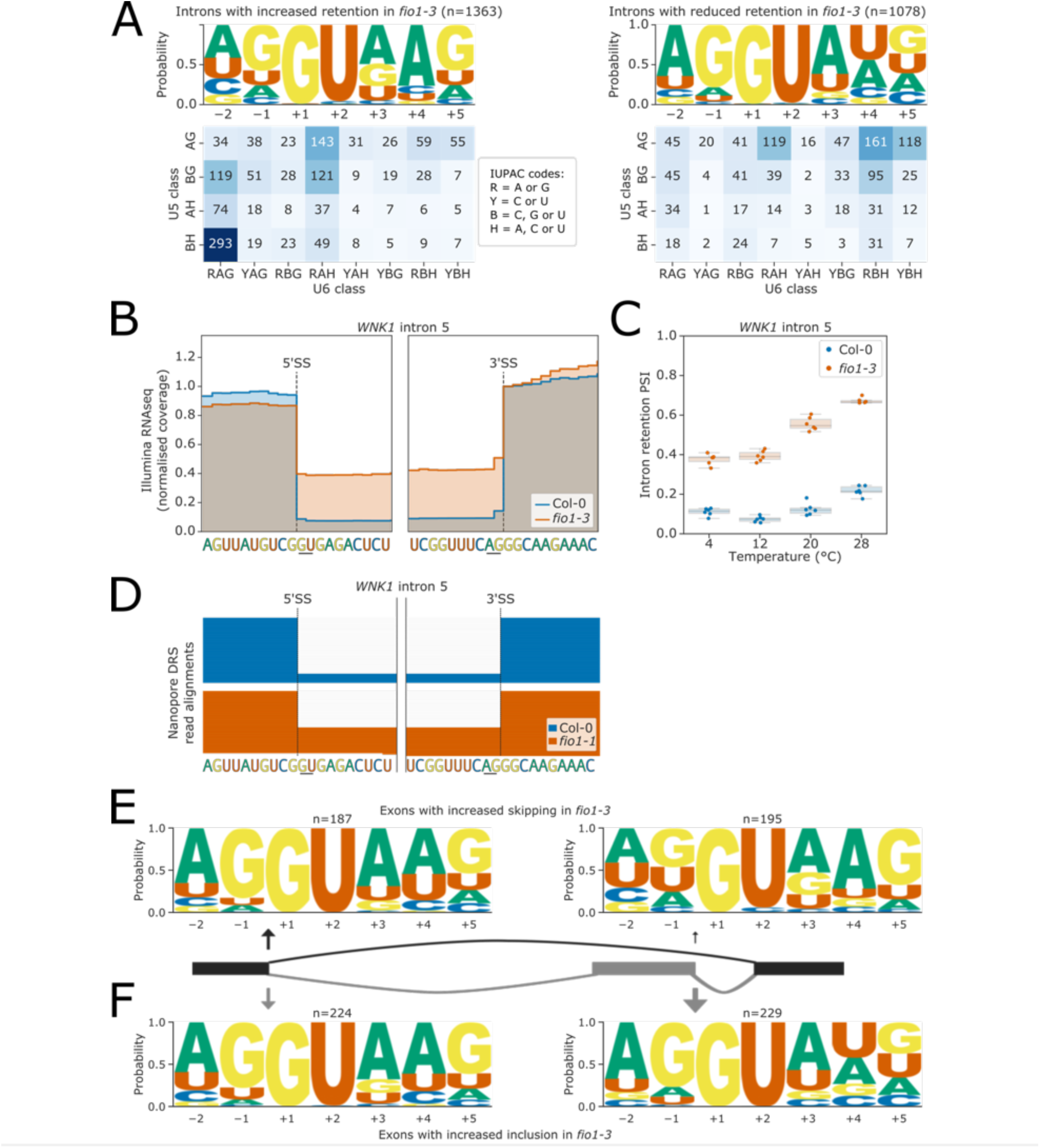
Effect of *fio1-3* on retained introns and exon skipping events. **(A)** Sequence logos and heatmaps showing the distribution of U5 snRNA and U6 snRNA interacting sequence classes for 5’SSs which have increased (left) and decreased (right) retention in *fio1-3*. Motifs are shown for the —2 to +5 positions of the 5’SS. U5 classes are based upon the distance of the —2 to —1 positions of the 5’SS from the consensus motif AG. U6 classes are based upon the distance of the +3 to +5 positions of the 5’SS from the consensus motif RAG. **(B)** Gene track showing intron retention at *WNK1* intron 5 in *fio1-3*, at 20°C. Expression is normalised by the coverage at the +1 position of the 3’SS. **(C)** Boxplot showing the change in intron retention of *WNK1* intron 5 at varying temperatures, in Col-0 and *fio1-3*. **(D)** Gene track showing intron retention at *WNK1* intron 5 in *fio1-1*, identified using nanopore DRS read alignments. Alignments have been subsampled to a maximum of 50 per condition. **(E-F)** Sequence logos for 5’SSs at introns upstream (left) and downstream (right) of exons with **(E)** increased skipping or **(F)** increased retention in *fio1-3*.

The remaining introns with significantly altered retention are more efficiently spliced in *fio1-3.* These introns tend to have stronger matches to U5 snRNA loop 1 in the 5’ exon, and only 34.7% have A_+4_, whilst 44.3% have U_+4_ (Figure 6A). This suggests that loss of m^6^A from U6 snRNA actually increases the splicing efficiency of introns with U at the +4 position of the 5’SS.

A common form of alternative splicing involves the excision of an exon and flanking introns in a splicing event called exon skipping. We identified 442 exons with significantly different levels of skipping in *fio1-3.* Of these, 53.6% have increased levels of inclusion and 46.4% have increased levels of skipping. We considered these two groups separately and performed 5’SS motif analysis.

We found that in cases where exon skipping increased in *fio1-3,* 71.3% of the 5’SSs at the downstream intron (i.e. at the 3’ end of the skipped exon) had A_+4_, in combination with weak U5 snRNA loop 1 interacting sequences (Figure 6E, Figure 6 – figure supplement 1A). In comparison, the 5’SS of the upstream intron was more likely to have a stronger U5 snRNA loop 1 interacting sequence (Figure 6E). This is consistent with a relative weakening of the recognition of the 5’SS at the downstream intron in *fio1-3*. An example of this phenomenon is illustrated by the gene encoding the floral repressor MAF3 (AT5G65060). At *MAF3*, there is an increase in skipping of exon 2 in *fio1-3* (Figure 6 – figure supplement 1B-C). The 5’SS at the 3’ end of this exon has the sequence UA//GUAAG, whereas the upstream 5’SS has the sequence AA//GUAAG, which is a stronger match to U5 snRNA loop 1.

In cases where exon inclusion increased in *fio1-3,* we found that 41.9% of 5’SSs at the 3’ end of the skipped exon had U_+4,_ compared to only 31.0% with A_+4_ (Figure 6F, Figure 6 – figure supplement 1A). In comparison, 60.7% of 5’SSs upstream of the skipped exon had A_+4_, indicating that the relative strength of the two 5’SSs changes in *fio1-3*. For example, at *PTB1* (AT3G01150), there is an increase in inclusion of exon 3 in *fio1-3* (Figure 6 – figure supplement 1D-E). The 5’SS at the 3’ end of this exon has the sequence AG//GUGUC, whereas the upstream 5’SS has the sequence UG//GUGAG. Although both the upstream and downstream 5’SSs are used when the exon is retained, it is possible that the relative strengthening of recognition of the downstream 5’SS in *fio1-3* improves exon definition.

In summary, we can identify and characterise changes in intron retention and exon skipping sensitive to loss of FIO1 function. Although the outcome of these splicing events is different from alternative 5’SS selection, they are associated with the same changes in 5’SS sequence as those associated with alternative 5’SS selection.

### Alternative 3’SS usage in *fio1-3* can be explained by U6 snRNA-dependent interactions between 5’ and 3’SSs

We observed 1484 instances where 3’SS selection was altered in *fio1-3*. In some cases, these alternative 3’SSs were linked with altered 5’SS choice or intron retention, but this accounted for a relatively small proportion of events (Figure 7 – figure supplement 1A). Alternative 3’SS selection was found to be equally likely to switch in an upstream or downstream direction in *fio1-3* (Figure 7A). There was a strong enrichment for very local switching of 3’SSs, with 37.9% of alternative 3’SSs occurring within 6 nt of the *fio1-3*- sensitive 3’SS, and 18.9% of examples occurring at exactly 3 nt upstream or downstream. These examples correspond to NAGNAG-like acceptors, which have previously been characterised in multiple species including human and Arabidopsis (Bradley et al., 2012; Hiller et al., 2004; Schindler et al., 2008).

**Figure 7.**
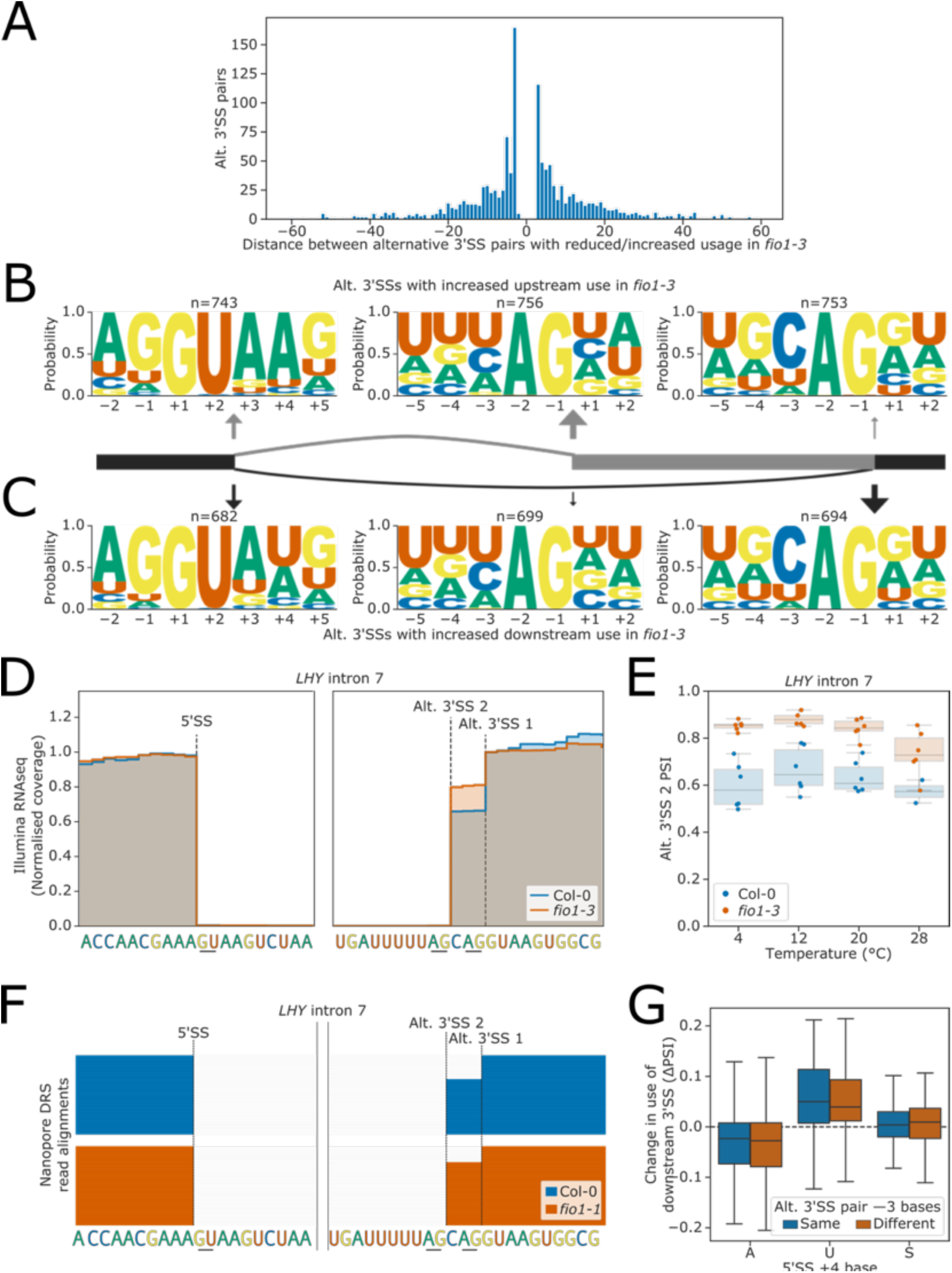
Effect of *fio1-3* on alternative 3’SS usage. **(A)** Histogram showing the distance between alternative 3’SS pairs with significantly different usage in *fio1-3*. Negative distances represent shifts towards greater usage of upstream 3’SSs, whilst positive distances represent shifts towards greater usage of downstream 3’SSs. **(B-C)** Sequence logos for 5’SSs (left), upstream 3’SSs (middle) and downstream 3’SSs (right) at pairs of alternative 3’SSs with increased **(B)** upstream or **(C)** downstream usage in *fio1-3.* 5’SS logos are for —2 to +5 positions, 3’SS logos are for —5 to +2 positions. **(D)** Gene track showing alternative 3’SS usage at *LHY* intron 5 in *fio1-3*, at 20°C. Expression is normalised by the coverage at the —1 position of the 5’SS. **(E)** Boxplot showing the change in usage of the upstream alternative 5’SS (Alt 5’SS 2) in *LHY* intron 5 at varying temperatures, in Col-0 and in *fio1-3*. **(F)** Gene track showing intron retention at *LHY* intron 5 in *fio1-1*, identified using nanopore DRS read alignments. Alignments have been subsampled to a maximum of 50 per condition. **(G)** Boxplot showing the change in usage of downstream 3’SSs in alternative 3’SS pairs with different 5’SS +4 bases, separated by whether the base at the —3 position of the two alternative 3’SSs is the same (e.g. CAG\\CAG\\) or different (e.g. UAG\\CAG\\).

To examine the 5’SSs associated with alternative 3’SSs used in *fio1-3*, we separated examples with increased upstream and downstream 3’SS usage and performed motif analysis. At introns with a relative increase in upstream 3’SS usage in *fio1-3*, we found that 80.5% of the corresponding 5’SSs were characterised by A_+4_ (Figure 7B). For example, at *LHY* (AT1G01060), we identified an AA//GUAAG 5’SS and a UAG\\CAG\\ alternative 3’SS pair in intron 7 (Figure 7D). In *fio1-3*, at 20°C, there is a 20.8% shift in favour of the upstream UAG\\ 3’SS (Figure 7E). This 3’SS switch is supported by orthogonal nanopore DRS analysis of the *fio1-1* allele (Figure 7F). Conversely, when we analysed the features of introns where a relative increase in downstream 3’SS usage was detected in *fio1-3,* we found that only 37.0% of these 5’SSs had A_+4_, whereas 48.8% had U_+4_ (Figure 7C). For example, at *MAF2* intron 4, we identified an AG//GUAUU 5’SS and a UAG\\ACAG\\ alternative 3’SS pair (Figure 7 – figure supplement 1B). In *fio1-3*, at 20°C, there is a 12.6% shift in favour of the downstream CAG\\ 3’SS (Figure 7 – figure supplement 1C).

3’SS choice involves scanning downstream from the branchpoint to the first available 3’SS motif (Smith et al., 1993). However, competition with downstream 3’SSs can occur within a short range, such as in NAGNAG acceptors. Downstream 3’SSs in *fio1*-sensitive alternative 3’SS pairs were more likely to have a cytosine as the —3 position than upstream 3’SSs (Figure 7B-C), a feature that has been shown to increase 3’SS competitiveness (Bradley et al., 2012; Smith et al., 1993). However, this appears to reflect differences in the background rate of C_-3_ at upstream and downstream 3’SSs, rather than a change in motif competitiveness in *fio1-3* (Figure 7 – figure supplement 1D). To analyse whether the —3 position contributes to changes in alternative 3’SS usage in *fio1*, we separated *fio1*-sensitive 3’SS pairs where the base at the —3 position was the same at both the upstream and downstream 3’SS, from examples which had different bases at the —3 position. We found that among both sets of 3’SS pairs, 5’SS A_+4_ was still associated with increased upstream usage in *fio1-3*, whereas U_+4_ was associated with increased downstream usage (Figure 7G). This demonstrates that changes in 3’SS usage in *fio1-3* are caused by a change in the competitiveness of distal 3’SSs, irrespective of 3’SS motif.

These findings link selection of the 3’SS and 5’SS. When U6 snRNA recognition of the 5’SS is favoured by either an m^6^A:A_+4_ interaction in WT Col-0 or A:U_+4_ interaction in *fio1-3,* this increases the usage of distal 3’SSs. When the U6 snRNA/5’SS interaction is less favoured, then upstream 3’SSs are more likely to be used. We conclude that the interactions of U6 snRNA with the 5’SS can influence usage of competing 3’SSs.

### Sequences targeted by U5 and U6 snRNAs are anticorrelated in the 5’SSs of developmentally complex eukaryotes

Our results indicate that a strong match to U5 snRNA loop 1 in the 3’ end of the upstream exon offsets the effects of 5’SS A_+4_ unfavourability in *fio1-3*. We therefore reasoned that if U5 and U6 cooperate in 5’SS selection, then strong U5 snRNA loop 1 interactions may globally compensate weaker U6 snRNA ACAGA box interactions, and *vice versa*. To test this hypothesis, we used all annotated Arabidopsis 5’SS sequences to generate position-specific scoring matrices (PSSMs) for the —2 to —1 positions corresponding to the U5 snRNA loop 1 interacting region, and the +3 to +5 positions corresponding to the U6 snRNA ACAGA box interacting region. We then used these PSSMs to score the U5 and U6 snRNA interaction log-likelihood of each individual 5’SS. 5’SSs with AG//GU motifs have high scoring U5 PSSM scores, and those with //GURAG motifs have high scoring U6 PSSM scores. This analysis revealed that U5 and U6 snRNA PSSM scores are indeed negatively correlated in Arabidopsis (Spearman’s ρ = —0.36, p < 1×10^—16^). We found that 5’SSs lacking a //GURAG motif had significantly higher U5 PSSM scores than //GURAG 5’SSs (Figure 8A, B), whereas BH//GU 5’SSs had significantly higher U6 PSSM scores than AG//GU 5’SSs (Figure 8C, D). We found similar anticorrelations in U5 and U6 PSSM scores in other species, including *C. elegans* (ρ = —0.38, p < 1×10^—16^; Figure 8 – figure supplement 1A), *Drosophila melanogaster* (ρ = —0.30, p < 1×10^—16^; Figure 8 – figure supplement 1B), *Danio rerio* (ρ = — 0.36, p < 1×10^—16^; Figure 8 – figure supplement 1C), and *Homo sapiens* (ρ = —0.17, p < 1×10^-16^; Figure 8 – figure supplement 1D). These findings indicate that there are two major classes of 5’SS in distinct eukaryotes: //GURAG and AG//GU. In the metazoan genomes that we examined, these two classes occur with approximately equal frequency (Figure 8 – figure supplement 1A-D). The combined stability of binding to U5 and U6 snRNA probably determines 5’SS selection after initial recognition by U1 snRNA and other protein factors. These findings indicate widespread compensatory differences in the relative strength of either U5 snRNA loop 1 or U6 ACAGA box interactions at 5’SSs.

**Figure 8.**
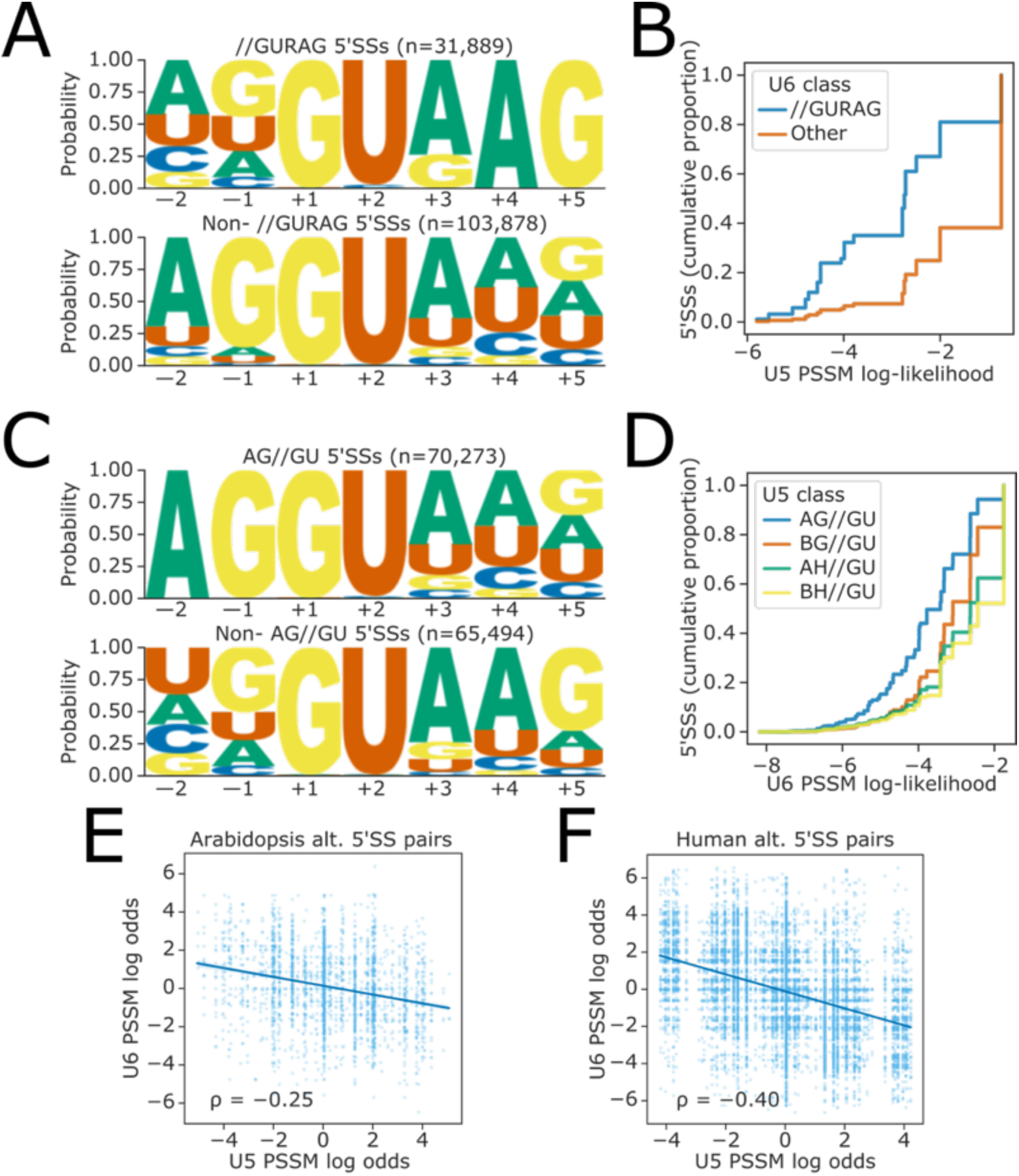
Global analysis of U5 and U6 interaction strengths reveals anticorrelation. **(A)** Sequence logos showing base frequency probabilities at —2 to +5 positions for (above) 5’SSs with //GURAG sequence and (below) all other 5’SSs. **(B)** Empirical cumulative distribution function of U5 PSSM log-likelihood scores for 5’SSs with either //GURAG sequence or all other 5’SSs. U5 PSSM scores are calculated using a PSSM derived from all 5’SSs in the Araport11 reference annotation, at the —2 to —1 positions of the 5’SS, inclusive. **(C)** Sequence logos showing base frequency probabilities at the —2 to +5 positions for (above) 5’SSs with AG//GU sequence and (below) all other 5’SSs. **(D)** Empirical cumulative distribution function of U6 PSSM log-likelihood scores for 5’SSs with different U5 classes. U6 PSSM scores are calculated using a PSSM derived from all 5’SSs in the Araport11 reference annotation, at the +3 to +5 positions of the 5’SS, inclusive. **(E-F)** Scatterplot showing the ratio of PSSM log-likelihoods (log-odds ratio) for U5 and U6 snRNA interacting sequences, at pairs of upstream and downstream alternative 5’SSs in **(E)** the Araport11 reference annotation or **(F)** the *H. sapiens* GRCh38 reference annotation. A positive log-odds ratio indicates that the PSSM score of the upstream 5’SS is greater than that of the downstream 5’SS.

### Opposing U5 and U6 snRNA interaction potential is a feature of alternative 5’SS pairs in developmentally complex eukaryotes

Given that many 5’SSs appear to have either strong U5 or U6 snRNA interacting sequences, we speculated that opposing U5 and U6 snRNA interaction strengths at pairs of alternative 5’SSs could contribute to alternative splicing. To test this hypothesis, we calculated the log-odds ratio of U5 and U6 PSSM scores for all pairs of annotated alternative 5’SSs in Arabidopsis. We found that these ratios were negatively correlated (Spearman’s ρ = —0.25, p < 1×10^—16^), indicating that 5’SS pairs do tend to have complementary strengths with respect to U5 and U6 snRNA recognition (Figure 8E). The strength of this complementary relationship varies in different organisms: in humans, we found a stronger negative correlation between relative U5 and U6 PSSM scores (ρ = —0.40, p < 1×10^—16^; Figure 8F), whereas in *C. elegans,* the correlation was weaker (ρ = —0.15, p = 8.61×10^—7^; Figure 8 – figure supplement 2A), and in *D. rerio*, there was no correlation (ρ = —0.01, p = 0.58; Figure 8 – figure supplement 2B). We conclude that changing the relative favourability of 5’SSs with stronger U5 or U6 snRNA interacting sequences, as occurs in *fio1-3*, could be a mechanism contributing to alternative 5’SS choice.

## Discussion

### FIO1 buffers splicing fidelity against temperature change

We identified *fio1* through a mutant screen designed to reveal factors that control the temperature-responsive splicing of mRNA encoding the floral repressor, MAF2. The *fio1* mutants have increased retention of *MAF2* intron 3 compared to WT Col-0. However, splicing of *MAF2* intron 3 remains responsive to temperature in *fio1* mutants, suggesting that FIO1 is not the thermosensor in this process. The level of *MAF2* intron 3 retention in *fio1* at 4°C and 20°C is similar to that detected in WT Col-0 at 12°C and 28°C, respectively, indicating that FIO1 function calibrates the temperature range over which *MAF2* alternative splicing occurs. Our global RNA-Seq analyses indicate that most splicing events disrupted in *fio1-3* were temperature-insensitive. However, we did detect a subset of changes to 5’SS selection in *fio1-3* (672 events) that became more extreme at elevated temperatures (e.g. 28°C). Therefore, FIO1 function ensures splicing fidelity by buffering against the impact of elevated temperatures.

### The splicing of mRNAs encoding regulators of flowering time is disrupted in *fio1*

All three *fio1* alleles studied here flower early. We found multiple splicing defects that disrupt the functional expression of not only MAF2, but also the floral repressors *FLM* and *MAF3* in *fio1* alleles. Detectable levels of sense and antisense RNAs at the locus encoding the floral repressor *FLC* were reduced in *fio1*, but no splicing changes were detected, suggesting that the impact of U6 snRNA m^6^A modification on *FLC* expression is indirect. FLM, MAF2, MAF3 and FLC function together with SVP in higher order protein complexes to repress the expression of *FT* and *SOC1.* We detected elevated transcript levels of *FT* and *SOC1* in *fio1* mutants, consistent with the idea that floral repressor activity had been compromised. We saw splicing changes in transcripts encoding circadian modulators such as *LHY* and *WNK1* (Kumar et al., 2011; Mizoguchi et al., 2002), which may contribute to the lengthening of the circadian period observed in *fio1* mutants (Kim et al., 2008). Finally, we found that aberrant splicing limits the functional expression of *AtSAR1* in *fio1* mutants. *AtSAR1* encodes a nucleoporin that shuttles the E3 ubiquitin ligase HOS1 to the nucleus, where it controls CO protein abundance (Dong et al., 2006; Li et al., 2020; Parry et al., 2006) and *FLC* gene expression (Jung et al., 2013). As a result, *hos1* and *sar1* mutants are early flowering due to increased CO protein levels (Li et al., 2020) and reduced *FLC* expression (Jung et al., 2013; Li et al., 2020). Both *hos1* and *sar1* mutants also have lengthened circadian periods (MacGregor et al., 2013). Overall, these findings suggest that the early flowering phenotype of *fio1* results from splicing changes that reduce the activity of floral repressors and increase the activity of factors that promote flowering, such as CO.

### FIO1-dependent m^6^A modification of U6 snRNA determines splicing accuracy and efficiency

Our data are consistent with the idea that the major effect of FIO1 occurs through m^6^A modification of U6 snRNA and the subsequent direct interaction of m^6^A modified U6 snRNA with target 5’SSs. Our nanopore DRS approach did not reveal widespread FIO1-dependent m^6^A sites in poly(A)+ mRNA. Consistent with this, our previous use of the orthogonal technique, miCLIP (which detects m^6^A sites covalently cross-linked to anti-m^6^A antibodies by UV light) mapped m^6^A in poly(A)+ RNA to mRNA 3’ terminal exons, but not splice sites or introns (Parker et al., 2020). We found that 8.7% of *FIP37-*dependent m^6^A sites also had slightly altered modification rates in *fio1-1,* suggesting that loss of *FIO1* might indirectly affect m^6^A sites written by the METTL3-like complex. Consistent with this idea, we detect defects in the splicing of pre-mRNA encoding the MTB (METTL14) subunit of this complex in *fio1* mutant alleles that may compromise its function. It is also possible that changes in patterns of gene expression caused by loss of FIO1 function perturb the canonical targeting of m^6^A in some transcripts by the METTL3-like writer complex. A combination of these phenomena, and different analytical approaches, might account for recent contrasting reports on FIO1-dependent mRNA methylation (Sun et al., 2022; Wang et al., 2022; Xu et al., 2022).

We do not rule out the possibility that FIO1 may have other RNA targets not detected by our approaches, but it is remarkable that we can classify the vast majority of *fio1*-sensitive splicing events simply based on sequence features of 5’SSs. When U6 snRNA is m^6^A modified, 5’SSs with A_+4_ are favoured. In the absence of U6 snRNA m^6^A, 5’SSs with U_+4_ and/or stronger interactions with U5 snRNA loop 1 are favoured. Cryo-electron microscopy (cryo-EM) structures of the human spliceosome suggest U6 snRNA m^6^A faces 5’SS_+4_ (Bertram et al., 2017). Consistent with this, we can separate almost 80% of FIO1-dependent 5’SS choices by the identity of the base at the +4 position of 5’SSs alone, underlining the probable direct effect of most of the splicing changes we detect. In comparison, no obvious difference in splice site sequences were associated with temperature-dependent alternative splicing.

Our findings are similar, in some respects, to those recently reported for *S. pombe*, where disruption of the FIO1 orthologue, MTL16, results in widespread intron retention (Ishigami et al., 2021). A more diverse set of splicing events is disrupted in Arabidopsis *fio1* mutants. Nevertheless, in both species, most changes in splicing can be explained by the impact of m^6^A modification on U6 snRNA recognition of the 5’SS_+4_ position, or the relative strength of U5 and U6 snRNA interactions at 5’SSs. Consistent with this, defective splicing of specific introns in *S. pombe mtl16*Δ strains can be experimentally rescued by expression of mutated U5 snRNAs designed to strengthen U5 loop 1 interactions with the upstream exon (Ishigami et al., 2021). Together, these findings reveal the impact of U6 snRNA m^6^A modification and indicate that U5 and U6 snRNAs have cooperative and compensatory roles in 5’SS selection dependent on 5’SS sequence (Figure 9A).

**Figure 9:**
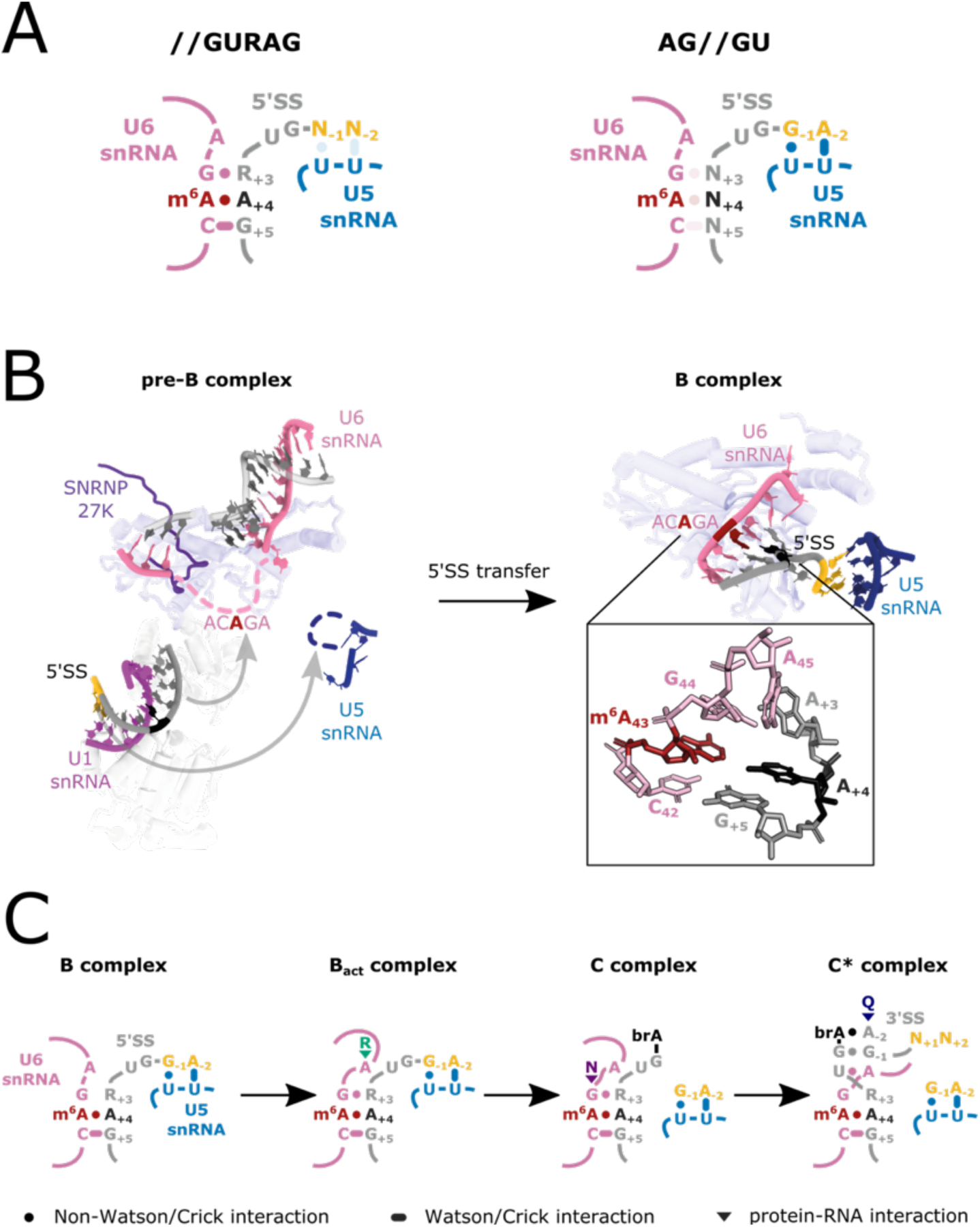
U6 m^6^A:5’SS_A+4_ interactions during splicing. **(A)** Model depicting U5 and U6 snRNA interactions with two major classes of 5’SS: //GURAG and AG//GU 5’SSs. //GURAG 5’SSs form strong interactions with U6 snRNA ACAGA (darkly shaded) and weaker interactions with U5 snRNA loop 1 (lightly shaded). AG//GU 5’SSs form strong interactions with U5 snRNA loop 1 and weaker interactions with U6 snRNA ACAGA. **(B)** Cryo-EM analysis of human pre-B and B complexes with RNA interactions detailed in the expanded section (PDB 6AHD) (Bertram et al., 2017) and Prp8 shown in the background as a common scaling reference. The U6 snRNA ACAGA and U5 snRNA loop 1 sequences are missing from cryo-EM structures at this stage, probably because they present as flexible loops. In B complex, C42 and 5’SS G_+5_ form a canonical Watson–Crick pair. m^6^A43 and 5’SS A_+4_ form a trans Hoogsteen sugar edge interaction (Leontis et al., 2002) that caps and stabilises the U6/5’SS helix by stacking because U6 snRNA G44 and 5’SS A_+3_ have not yet formed a stable interaction. The 5’SS is kinked and U5 snRNA loop 1 is docked on the upstream exon. The methyl group of U6 snRNA m^6^A43 is not modelled in the structure due to lack of resolution. **(C)** Model depicting U6 m^6^A interactions at different stages of splicing. In B complex, U6 m^6^A43 stabilises the U6/5’SS helix by stacking. As the active site forms in B_act,_ this role becomes less important because U6 snRNA G44 interacts more stably with 5’SS_+3_ and U6 A45 stacks on the helix stabilised by R554 of SF3B2. In C complex, the U6/5’SS helix is stabilised by N57 of hYJU2. In C* complex, the U6m^6^A43:5’SS_+4_ interaction becomes more important again because the 5’SS_+3_ pivots to a new position. The m^6^A43 and 5’SS_+4_ pair forms part of a continuous helical stack with the docked 3’SS, which is capped by the interaction between 3’SS_-3_ and Q1522 of Prp8. For more detail, see Figure 9 – supplement 1.

### Linking cooperative and compensatory roles for U5 and U6 snRNA in 5’SS selection, and the evolution of two major classes of 5’SS

Our global analyses of annotated Arabidopsis 5’SSs identified anti-correlated biases in sequence composition at U5 and U6 snRNA interacting positions. We found the same to be true for 5’SSs in other eukaryotes, including humans. Published 5’SS consensus motifs typically aggregate all 5’SSs. However, on careful analysis two different classes of human 5’SS motifs emerge: //GURAG, which accounts for 54% of 5’SSs, and AG//GU, which accounts for 45% of 5’SSs (Sibley et al., 2016), indicating that this basic feature of human genome organization has been hiding in plain sight. Compensatory patterns of base composition at human 5’SSs have been described but not explained (Burge and Karlin, 1997; Carmel et al., 2004; Wong et al., 2018). For example, a “seesaw linkage” pattern was observed where −1G permits any nucleotide at position +5 and conversely, +5G permits any nucleotide at position −1 (Burge and Karlin, 1997). Our findings suggest that such compensatory base composition at 5’SSs can be accounted for by two major classes of 5’SS recognised mainly by interactions with either U5 snRNA loop 1 or the U6 snRNA AC**<u>A</u>**GA box. Furthermore, the cooperativity of U5 and U6 snRNA function in 5’SS selection may explain why having Gs at both −1 and +5 positions is highly preferential for efficient splicing (Wong et al., 2018). We could extend this finding to other eukaryotes, including *C. elegans*, *D. melanogaster* and *D. rerio*. Compensatory interactions are likely to facilitate degeneracy in sequences that can be recognised as 5’SSs. Degeneracy may act as a buffer against deleterious mutations in splice site sequences, as well as lowering barriers to the evolution of new splicing structures. This is important because alternative splicing is clearly associated with developmental complexity (Bush et al., 2017; Nilsen and Graveley, 2010).

Alternative splicing is the result of competition between multiple splicing choices. When we analysed annotated alternative 5’SS pairs in Arabidopsis, we found that one of the alternative 5’SS pair was more likely to exhibit strong U5 snRNA loop 1 recognition, and the other had stronger matches to U6 snRNA AC**<u>A</u>**GA box. We also found that in the absence of U6 snRNA m^6^A modification the splicing of introns with 5’SS U_+4_ became more efficient. This suggests that alternative splicing could be regulated by changing the relative favourability of U5 and U6 snRNA interactions. The extent of opposing U5 and U6 snRNA interactions at alternative 5’SSs varied across eukaryotes, suggesting that this form of regulation may only be used in specific organisms.

The simplification of splicing in *S. cerevisiae* may reflect an extreme outcome of the evolution of compensatory roles for U5 and U6 snRNA in 5’SS choice because the U6 snRNA ACAGA interacting positions of *S. cerevisiae* 5’SSs are almost completely invariant, whereas the U5 snRNA loop 1 interacting positions in the upstream exon are completely degenerate (Neuvéglise et al., 2011). Given that *S. cerevisiae* almost completely lacks alternative splicing, it may be that varying U5/U6 interaction strengths is no longer required, and so a preference for strong U6 snRNA ACAGA interacting positions removes the selective pressure on U5 snRNA loop 1 interacting positions in protein coding exons.

Since m^6^A modified U6 snRNA contributes to the selection of 5’SSs that are less constrained by upstream exon sequence composition, more degenerate sites can be selected. It is therefore possible that the evolution of alternative splicing is associated with U6 snRNA m^6^A modification, cooperative and compensatory roles for U5 and U6 snRNA and the evolution of spliceosomal proteins that stabilise and proofread these RNA interactions.

### What is the role of the m^6^A modification in U6 snRNA?

Our analysis of available cryo-EM structures of human spliceosomes (Bertram et al., 2017, 2020; Fica et al., 2019; Zhang et al., 2018) does not provide clear evidence of a direct interaction between U6 snRNA m^6^A and a spliceosomal protein (Figure 9B; Figure 9 – figure supplement 1). Instead, during the transfer of the 5’SS from U1 snRNA to U6 and U5 snRNAs in spliceosomal B complexes, U6 AC^m6^**<u>A</u>**GA m^6^A faces 5’SS A_+4_ in a trans Hoogsteen sugar edge interaction that could stabilise the 5’SS/U6 helix by capping (Figure 9B). This is important because 5’SS A_+3_ is not aligned to stack on A_+4_ at this stage. However, later in spliceosomal B^act^ and C complexes, 5’SS A_+3_ engages in a more stable interaction with the U6 snRNA G44 adjacent to m^6^A (AC^m6^A<u>G</u>A), stacking on top of the 5’SS/U6 helix and stabilising it (Figure 9C; Figure 9 – figure supplement 1). Such stabilisation is important because degenerate 5’SS sequences in developmentally complex eukaryotes, mean that the helix formed with U6 snRNA AC**A**GA is relatively weak, short, and comprised mostly of non-canonical RNA-RNA interactions (Figure 9B).

Our global RNA-Seq analyses reveal that m^6^A-modified U6 snRNA rarely selects 5’SS with U_+4_, preferring instead to pair with 5’SSs with A_+4_. Biophysical data from model RNAs provide a possible explanation for this finding: an m^6^A-U base pair can form in a duplex, but the methylamino group rotates from its energetically preferred *syn* geometry on the Watson–Crick face to the higher-energy *anti* conformation, positioning the methyl group in the major groove (Roost et al., 2015). As a result, m^6^A has a destabilising effect on A-U base-pairs in short RNA helices (Kierzek and Kierzek, 2003; Roost et al., 2015). Therefore, 5’SS U_+4_ may be selected less frequently by m^6^A-modified U6 snRNA because of destabilisation of an already weak helix. In contrast, the thermal stability of m^6^A:A is increased compared to A:A (Roost et al., 2015).

Biophysical measurements also indicate m^6^A in unpaired positions base stacks more strongly than the unmodified base, adding substantial stabilization to adjacent duplexes (Roost et al., 2015). Consistent with this observation, global mapping of the structural context around m^6^A shows a general transition from single-stranded to double stranded regions around m^6^A sites (Roost et al., 2015). The non-canonical trans Hoogsteen sugar edge interaction between U6 snRNA m^6^A and 5’SS A_+4_, detected in cryo-EM analysis of splicesome B complexes, also suggests the methyl group could stabilise the U6/5’SS helix by stacking (Bertram et al., 2017). Such helix capping by m^6^A that does not rely on a very strong Watson–Crick interaction may favour a single-stranded conformation of the adjacent 5’SS_+1-3_, which is needed for a kink in the RNA to form. These structural changes in B complex enable the upstream exon to dock onto U5 snRNA loop 1 and align the 5’SS cleavage site in preparation for formation of the active site. In the absence of U6 snRNA m^6^A modification, our data reveal that 5’SS U_+4_ sites are preferred. This is consistent with a relatively strong interaction being important at the terminal position of the U6 snRNA ACAGA helix in B complex to stabilise its formation. However, we found that this was not necessarily sufficient for 5’SS selection because switches in 5’SS use were often accompanied by stronger interaction potential between U5 snRNA loop 1 and the upstream exon. These observations are consistent with a concerted process for transfer of the 5’SS from the U1 snRNP to U6 and U5 snRNAs during the transition from the pre-B to the B complex (Charenton et al., 2019, Figure 9-figure supplement 1).

The potential role of U6 snRNA m^6^A in 3’SS selection may be explained by U6 AC**<u>A</u>**GA interactions during remodelling of the spliceosome for the second splicing reaction (Figure 9C; Figure 9 – figure supplement 1). In humans, RNA rearrangements during the C to C* transition include stacking of the 3’SS G_-1_ onto U6 A45 (AC**A**G<u>A</u>), which remains paired to U_+2_ of the 5’SS (Fica et al., 2019, 2017; Wilkinson et al., 2017). This results in a continuous helix stack, involving the interaction between U6 A43 and 5’SS A_+4_, that forms the receptor onto which the 3’SS docks. Our data are consistent with a model in which a strong interaction between U6 and the 5’SS +4 position stabilises this receptor, enabling 3’SSs distal to the BP to compete more efficiently with proximal 3’SSs that are favoured by scanning (Smith et al., 1993). A more stable receptor could allow the ATPase Prp22, which proof reads exon ligation (Mayas et al., 2006), to promote more efficient sampling and usage of distal 3’SSs. This has previously been observed in yeast (Semlow et al., 2016), where the U6/5’SS helix is intrinsically stronger than in Arabidopsis and humans (Plaschka et al., 2019; Wilkinson et al., 2020).

In summary, we suggest that the role of m^6^A in U6 snRNA is to stabilise the weak helix formed with the 5’SS and possibly to influence local RNA geometry. This role may explain the buffering function we discovered for FIO1 in 5’SS selection at elevated temperatures.

### A regulatory or adaptive role for U6 snRNA m^6^A modification?

It is an open question as to whether m^6^A modification of U6 snRNA is regulated. Either the activity of FIO1 might be controlled, or demethylases might act directly upon U6 snRNA. There is a precedent for control of splicing by regulation of U6 snRNA modifications because pseudouridylation of *S. cerevisiae* U6 snRNA, which controls entry into filamentous growth, can increase splicing efficiency of suboptimal introns (Basak and Query, 2014). Human UsnRNAs also appear to be targeted by demethylases. For example, FTO targets cap adjacent methylation (m^6^Am) in UsnRNAs (except U6) and this activity may account for splicing differences detected in FTO knockout backgrounds (Mauer et al., 2019).

It is also possible that natural genetic variation in *FIO1* is adaptive because two genome wide association studies have implicated variation at or near the *FIO1* locus in altered flowering time (Price et al., 2020; Sasaki et al., 2015). Consistent with our findings on flowering repressors, the impact of this variation on flowering time was the same at 10°C and 16°C. It will be interesting to test whether genetic variation alters the efficiency of U6 snRNA m^6^A modification by FIO1 in different ecotypes, and what impact this has upon global splicing patterns, including for the regulators of flowering time that we characterise here. There are 13 genes encoding U6 snRNA in the Arabidopsis Col-0 genome (Wang and Brendel, 2004). Currently, we know little about the relative patterns of U6 snRNA gene expression or m^6^A modification status, and how this may vary in different ecotypes. Indeed, this paucity of knowledge on U6 snRNA variants’ modification and expression applies to humans too. Our study on U6 snRNA m^6^A modification has important implications for understanding the mechanism of splicing and evolution of alternative splicing. It will now be interesting to investigate the possibility that modulation of U6 snRNA m^6^A may be regulatory or adaptive.

## Materials and methods

### Plant material

The wild type Col-0 accession, *fio1-3* (SALK_084201) and *maf2* (SALK_045623) were obtained from Nottingham Arabidopsis Stock Centre. The *fio1-1* mutant was a gift from Prof. Hong Gil Nam (Daegu Gyeongbuk Institute of Science and Technology), Republic of Korea. The *fip37-4* mutant was a gift from Prof. Rupert Fray (University of Nottingham), UK.

### Plant Growth Conditions

Seeds of wild-type Col-0, *fip37-4* and *fio1-1* used for nanopore DRS and m^6^A immunopurification were surface-sterilised and sown on MS10 media plates supplemented with 2% agar, stratified at 4°C for two days, germinated in a controlled environment at 20°C under 16 h light/8 h dark conditions, and harvested 14 days after transfer to 20°C. Seeds of wild-type Col-0 and *fio1-3* used for Illumina RNA-seq were surface sterilised and sown on 0.5 MS media plates supplemented with 2% agar, stratified at 4°C for two days, germinated in a controlled environment at 20°C under 16 h light/8 h dark conditions for 8 days. Seedlings were then transferred to either 28°C, 20°C, 12°C or 4°C, for 4 h under dark conditions before harvesting. Seeds used for *MAF2* splicing analysis were sown on 0.5 MS media plates supplemented with 2% agar, stratified at 4°C for two days, germinated in a controlled environment at 20°C under 16 h light/8 h dark conditions, and harvested 14 days after transfer to 20°C.

### Mutant screen

The two-step EMS mutant screen was conducted in a Col-0 line carrying a homozygous transgene in which the genomic *MAF2* coding region was translationally fused to luciferase (*gMAF2:LUC*). A Gateway cloning approach was used to introduce *gMAF2:LUC* into the Alligator vector pFP101, which features a recombination site downstream of a CaMV 35S promoter and selection by GFP fluorescence (Bensmihen et al., 2004). *Agrobacterium tumefaciens* strain GV3101 was used to transform Col-0 via the floral dip method (Clough and Bent, 1998). A homozygous line with strong LUC expression and a clear GFP fluorescence in the seed coat (Alligator selective marker) was identified and used for EMS mutagenesis. Approximately 20,000 M0 seeds of the *gMAF2:LUC* line were soaked overnight in 100 mL of phosphate buffer at 4°C. The buffer was replaced, with EMS added to a final concentration of 25 mM and the seeds were incubated at room temperature with gentle agitation for 16 h. The EMS was neutralised with 1M NaOH and the seeds were gently washed twice with 100 mM Sodium thiosulphate for 15 min, followed by three 15 min washes in distilled water. The M1 seeds were air dried on filter paper overnight, before planting on soil. The resulting plants were allowed to self-pollinate and grow to maturity. M2 seeds were collected from all plants and approximately 15,000 seeds were screened in two steps. In step 1, stratified seeds were planted on soil and grown in controlled environment chambers at 16°C, 16 h light/8 h dark, with light intensity of 200 µmol m^-2^ s^-1^. The first 100 plants to flower were selected for further screening. Splicing of *MAF2* intron 3 was assessed by RT-qPCR to identify early flowering lines showing enhanced intron retention at 16°C. Leaf material was frozen and homogenized using QIAGEN TissueLyser LT. RNA was extracted using a Nucleospin II RNA extraction kit (Machery–Nagel). Total RNA (1.5 µg per sample) was reverse-transcribed using the High Capacity cDNA Reverse Transcription Kit with Rnase Inhibitor (Applied Biosystems). Quantitative real time PCR was performed using GoTaq Hot Start Polymerase (Promega). Amplification of *MAF2* was performed according to manufacturer’s guidelines using primers 3 and 4 from (Rosloski et al., 2013), for 35 PCR cycles. These primers, which span *MAF2* intron 3, generate two products, which correspond to *MAF2 var2* (*MAF2* intron 3 retained) and *MAF2 var1* (*MAF2* intron 3 excised), respectively. PCR amplified products were separated on a 2% w/v agarose gel to resolve the splice variants. M2 plants showing a significantly enhanced *var2:var1* ratio compared to the parental line control were selected for further testing. The causative mutation was identified by bulked segregant analysis, sequencing pooled genomic DNA from phenotypically early flowering and wild type flowering time plants in a segregating F2 population. Genomic DNA was extracted using a Qiagen Plant DNA Maxi kit. Sequencing was carried out by the University of Leeds NGS facility using a HiSeq3000 sequencer with a 150 bp paired end library. The resulting DNA sequences were analysed using the artMAP pipeline (Javorka et al., 2019) unambiguously identifying a small region of chromosome 2 that is linked to the phenotype. Four genes in this region showed polymorphisms co-segregating with the phenotype: AT2G16770, AT2G19430, AT2G19720 and AT2G21070. For the first three of these candidates the SNP was in the 5’ UTR and caused a synonymous mutation, or fell in the middle of the first intron, respectively. In the case of AT2G21070, the identified mutation is a G to A change at position 9041454 that disrupts the conserved GT of the splice donor for intron 2.

### Nanopore DRS

#### Total RNA isolation

Total RNA was isolated using Rneasy Plant Mini kit (Qiagen) and treated with TURBO Dnase (ThermoFisher Scientific). The total RNA concentration was measured using a Qubit 1.0 Fluorometer and Qubit RNA BR Assay Kit (ThermoFisher Scientific). RNA quality and integrity was assessed using the NanoDrop™ 2000 spectrophotometer (ThermoFisher Scientific) and Agilent 2200 TapeStation System (Agilent).

#### Preparation of libraries for direct RNA sequencing of poly(A)+ mRNA using nanopores

Total RNA was isolated from the Col-0, *fip37-4* and *fio1-1* seedlings as detailed above. mRNA isolation and preparation of nanopore DRS libraries (using the SQK-RNA002 nanopore DRS Kit (Oxford Nanopore Technologies) were performed as previously described (Parker et al., 2020). Libraries were loaded onto R9.4 SpotON Flow Cells (Oxford Nanopore Technologies) and sequenced using a 48 h runtime. Four biological replicates were performed for each genotype.

#### Nanopore DRS mapping

Nanopore DRS reads were basecalled using Guppy version 3.6.0 high accuracy RNA model. For mRNA modification analysis, reads were mapped to the Araport11 reference transcriptome (Cheng et al., 2017) using minimap2 version 2.17 (Li, 2018), with parameters -a -L –cs=short -k14 –for-only – secondary=no. For other analyses, reads were mapped to the Arabidopsis *TAIR10* genome (Arabidopsis Genome Initiative, 2000) using two pass alignment with minimap2 version 2.17 and 2passtools version 0.3 (Parker et al., 2021b). First pass minimap2 alignment was performed using the parameters -a -L –cs=short -x splice -G20000 –end-seed-pen 12 -uf. 2passtools score was then run with default parameters on each replicate to generate junctions, followed by 2passtools merge to combine them into a final set of guide junctions. Reads were remapped with minimap2 using the same parameters, but with the addition of the guide junctions using –-junc-bed and –junc-bonus=10. Pipelines for processing of nanopore DRS data were built and executed using Snakemake version 6.15.3 (Köster and Rahmann, 2012).

#### Nanopore poly(A)+ mRNA modification analysis

Differential modification analysis was performed on Col-0, *fip37-4,* and *fio1-1* data using the “n- sample” GitHub branch of Yanocomp (Parker et al., 2021a). Kmer-level signal data was generated using f5c eventalign version 0.13.2 (Gamaarachchi et al., 2020; Loman et al., 2015) and Yanocomp prep. A three-way comparison between the genotypes was performed using Yanocomp gmmtest, with a minimum KS statistic of 0.25. A 5% false discovery rate threshold was used to identify transcriptomic sites with significant changes in modification rate. Motif analysis was performed using meme version 5.1.1 with the parameters -cons NNANN -minw 5 -maxw 5 -mod oops (Bailey et al., 2015). Differential poly(A) site usage was performed using d3pendr version 0.1 with default parameters, and thresholded using a 5% false discovery rate and an effect size (measured using earth mover distance) of 25 (Parker et al., 2021c).

### Illumina RNA sequencing

#### Preparation of libraries for Illumina RNA sequencing

Total RNA was isolated from Col-0 and *fio1-3* seedlings using Nucleospin RNA kit (Macherey–Nagel, 740955) and treated with rDNase (Macherey–Nagel, CAS 9003-98-9) on columns according to manufacturers’ instructions. RNA concentration, quality and integrity was assessed using the NanoDrop® 1000 spectrophotometer (Labtech) and agarose gel electrophoresis. Poly(A)+ RNA purification and Illumina RNA-Seq library preparation was performed by Genewiz UK Ltd. Poly(A)+ RNA was selected with NEBNext Poly(A) mRNA Magnetic Isolation Module. Preparation of the sequencing libraries was performed using the NEBNext Ultra II Directional RNA Library Prep Kit for Illumina (New England Biolabs). 150-bp paired-end sequencing was carried out using Illumina Novaseq 6000. Six biological replicates were performed for each genotype.

#### Illumina RNA sequencing data processing

Illumina RNA-Seq data was assessed for quality using FastQC version 0.11.9 and MultiQC version 1.8 (Andrews, 2017; Ewels et al., 2016). Reads were mapped to the TAIR10 genome using STAR version 2.7.3a (Dobin et al., 2013) with a splice junction database generated from the Araport11 reference annotation (Cheng et al., 2017). We used Stringtie version 2.1.7 in mix mode (Kovaka et al., 2019) to generate *de novo* transcriptome assemblies from Illumina RNAseq replicates and pooled Nanopore DRS data. All assemblies were merged with the Araport11 reference annotation using Stringtie merge to create a unified set of transcripts for quantification. Transcript open reading frames were annotated using Transuite version 0.2.2 (Entizne et al., 2020), which was also used to predict nonsense mediated decay sensitivity. Pipelines were written and executed using Snakemake version 6.15.3 (Köster and Rahmann, 2012).

Transcripts were quantified using Salmon version 1.1.0 (Patro et al., 2017) with the TAIR10 genome assembly as decoys. SUPPA version 2.3 was used to generate event level percent spliced indices (PSIs) from transcript level quantifications (Trincado et al., 2018). PSIs were loaded into Python 3.6.7 using pandas version 1.0, and generalised linear models were fitted per event using statsmodels version 0.11 (Harris et al., 2020; McKinney, 2010; Oliphant, 2007; Seabold and Perktold, 2010). GLMs were used to test the relationship of PSI with genotype, temperature, and genotype × temperature interaction. Calculated p-values were adjusted for multiple testing using the Benjamini-Hochberg false discovery rate method. A false discovery rate of 5% was chosen to threshold events with significant changes in PSI which correlated with genotype, temperature, or genotype × temperature. Changes in motif composition were tested using G-tests of base frequencies at the —2 to +5 position of the splice site. To generate sequence logos, 5’ or 3’ splice sites from sets of alternative splicing events were filtered to remove duplicated positions, and probability logos were plotted using matplotlib version 3.3 and matplotlib_logo (Hunter, 2007; Parker, n.d.). Contingency tables of splice site classes at U5 and U6 interacting positions were generated using the difference of the —2 to —1 positions of the 5’SS from the consensus motif AG, and the difference of the +3 to +5 positions of the 5’SS from the consensus motif RAG, respectively. Classes were ordered by their log-likelihoods using PSSMs generated from all annotated 5’SSs (described below). Heatmaps of contingency tables were generated with seaborn version 0.11 (Waskom, 2021). Gene tracks using reads aligned to the TAIR10 reference genome were generated using pyBigWig version 0.3.17, pysam version 0.18, and matplotlib version 3.3 (Heger et al., 2014; Hunter, 2007; Ramírez et al., 2014).

#### Global splice site analyses

To measure the predicted strengths of U5 snRNA loop 1 and U6 snRNA ACAGA box interactions for individual sequences, all 5’SSs annotated in the Araport11 reference transcriptome (Cheng et al., 2017) were used to generate log transformed position specific scoring matrices (PSSMs). The U5 and U6 log-likelihood scores for individual sequences were then calculated using the —2 to —1 and +3 to +5 positions, respectively. Correlation of U5 and U6 PSSM scores was calculated using Spearman rank correlation coefficient. This analysis was repeated using 5’SSs from the *Homo sapiens* GRCh38 (International Human Genome Sequencing Consortium, 2004), *Caenorhabditis elegans* Wbcel235 (The C. elegans Sequencing Consortium, 1998), *Drosophila melanogaster* BDGP6 (dos Santos et al., 2015), and *Danio rerio* GRCz11/danRer11 (Howe et al., 2013) genome assemblies and annotations, downloaded from Ensembl Genomes release 104 (Howe et al., 2021). To measure relative U5 and U6 interactions for pairs of alternative 5’SSs, SUPPA was used to identify pairs of alternative 5’SSs in each genome annotation (Trincado et al., 2018). These alternative 5’SS pairs were ordered by their genomic positions relative to the strand of the parent gene (i.e. upstream and downstream), and log-odds ratios were calculated, using downstream 5’SS log-likelihoods as the denominator. Correlation of U5 and U6 log-odds ratios was calculated using Spearman rank correlation coefficient. Scatter plots with linear regression lines were plotted using seaborn (Waskom, 2021), and 95% confidence intervals for regression lines were calculated using bootstrap sampling with replacement.

### Immunopurification and detection of m^6^A modified RNA

#### Synaptic Systems anti-m^6^A

Total RNA was purified from ∼300 mg of frozen plant tissue using the miRVana miRNA isolation kit (Ambion) and treated with DNase I (New England Biolabs), according to the manufacturer’s instructions. The quantity and integrity of RNA was checked using a NanoDrop 2000 spectrophotometer (Thermo Fisher Scientific) and Agilent 2200 TapeStation System. Approximately 5 μg of RNA was suspended in 500 μL low salt buffer (50 mM Tris-HCl pH 7.4, 150 mM NaCl, 0.5% v/v NP-40), with 80U RNAsin Plus RNase inhibitor, (Promega) and 10 μL (1mg/mL) m^6^A-rabbit polyclonal purified antibody (#202 003 Synaptic Systems). The samples were mixed by rotation for 2 h at 4°C. Protein A/G magnetic beads (Pierce™, ThermoFisher) were washed twice with low salt buffer, then added to each RNA/antibody sample and mixed with rotation at 4°C for 16 h. The beads were washed twice with high salt buffer (50 mM Tris-HCl pH 7.4, 0.5 M NaCl, 1% v/v NP-40), twice with low salt buffer, and twice with PNK wash buffer (20 mM Tris-HCl pH 7.4, 10 mM MgCl2, 0.2% v/v Tween-20). Immunopurified RNA was eluted by digestion with proteinase K in 200 μL of PK buffer (50 mM Tris-HCl pH 7.4, 150 mM NaCl, 1 mM EDTA, 0.1% w/v SDS) at 37°C for 20 min with shaking at 300 rpm, followed by phenol-chloroform extraction and sodium acetate/ethanol precipitation.

Immunopurifed RNA (45 ng) was reverse transcribed (RT) with U6 and U2 snRNA reverse primers using SuperScript™ III Reverse Transcriptase (ThermoFisher), according to the manufacturer’s instructions. RT-qPCR was carried out using the SYBR Green I (Qiagen) mix with primers targeted to U2 and U6 snRNA (Supplementary file 2). The specificity of RT-qPCR was confirmed by sequencing of amplified products.

#### Millipore anti-m^6^A

Total RNA was purified from ∼600 mg of frozen plant tissue using the miRVana miRNA isolation kit (Ambion) and treated with DNase I (New England Biolabs), according to the manufacturer’s instructions. The quantity and integrity of RNA was checked using a NanoDrop 1000 spectrophotometer (Thermo Fisher Scientific). After sodium acetate/ethanol precipitation approximately 10 μg of RNA was resuspended in 20 μL nuclease free water, with 5% of the RNA being kept for the input sample.

Protein A/G magnetic beads (Millipore, 16-663) were washed twice with m^6^A wash buffer (10mM TrisHcl pH7.4, 150mM NaCl, 0.1% v/v NP-40), and then resuspended in 50 μL m^6^A wash buffer. 5 μL (1mg/mL) of Anti-N6-methyladenosine (m^6^A) antibody (Millipore, ABE572), or Rabbit Anti-IgG (Thermofisher, A16104) was coupled to the washed beads on a roller for 40 mins at room temperature. After incubation, beads were washed three times in m^6^A wash buffer. m^6^A immunoprecipitation buffer was then added to the beads (10mM TrisHcl pH7.4, 150mM NaCl, 0.1% v/v NP-40, 17.5mM EDTA pH 8) with 80U SUPERaseIn™ RNase Inhibitor, and 10 μL of each RNA sample was added to both the anti-m6A and anti-IgG antibody/beads mixture. Samples were mixed by rotation at 4°C for 16 h. The beads were washed five times with m^6^A wash buffer. Immunopurified and 5% input RNA was eluted by digestion with proteinase K in 150 μL of PK buffer (50 mM Tris-HCl pH 7.4, 150 mM NaCl, 0.1%v/v NP-40, 0.1% w/v SDS) at 55°C for 30 minutes, followed by extraction with TRIzol™ LS (10296028, Thermofisher) and sodium acetate/ethanol precipitation.

Immunopurifed RNA (500 ng) was reverse transcribed (RT) with random hexamers using ThermoFisher MultiscribeTM II Reverse Transcriptase (4311235, Thermo Fisher Scientific) according to the manufacturer’s instructions. RT-qPCR was carried out using the SYBR Green I (Qiagen) mix with primers targeted to U2 and U6 snRNA (Supplementary file 2). The specificity of RT-qPCR was confirmed by sequencing of amplified products.

### Code availability

All pipelines, scripts and notebooks used to generate figures are available from GitHub at https://github.com/bartongroup/Simpson_Davies_Barton_U6_methylation and Zenodo at https://zenodo.org/record/6372644 (Parker, 2022).

### Data availability

Illumina sequencing data from the genetic screen that identified *fio1-4* is available from ENA accession PRJEB51468. Col-0, *fip37-4* and *fio1-1* nanopore DRS data is available from ENA accession PRJEB51364. Col-0 and *fio1-3* Illumina RNA-Seq data is available from ENA accession PRJEB51363.

## Acknowledgments

We thank Mary McKay (School of Biology, University of Leeds) for excellent technical assistance with the EMS mutant screen. We are indebted to Oliver Manners for help in establishing m^6^A immunoprecipitation. We thank David Lilley, Tim Wilson and Adrian Whitehouse for helpful discussions. We thank Alper Akay, Martin Balcerowicz, James Lloyd and Anjil Srivastava for comments on the manuscript.

## FIGURE SUPPLEMENTS

**Figure 3 – figure supplement 1.**
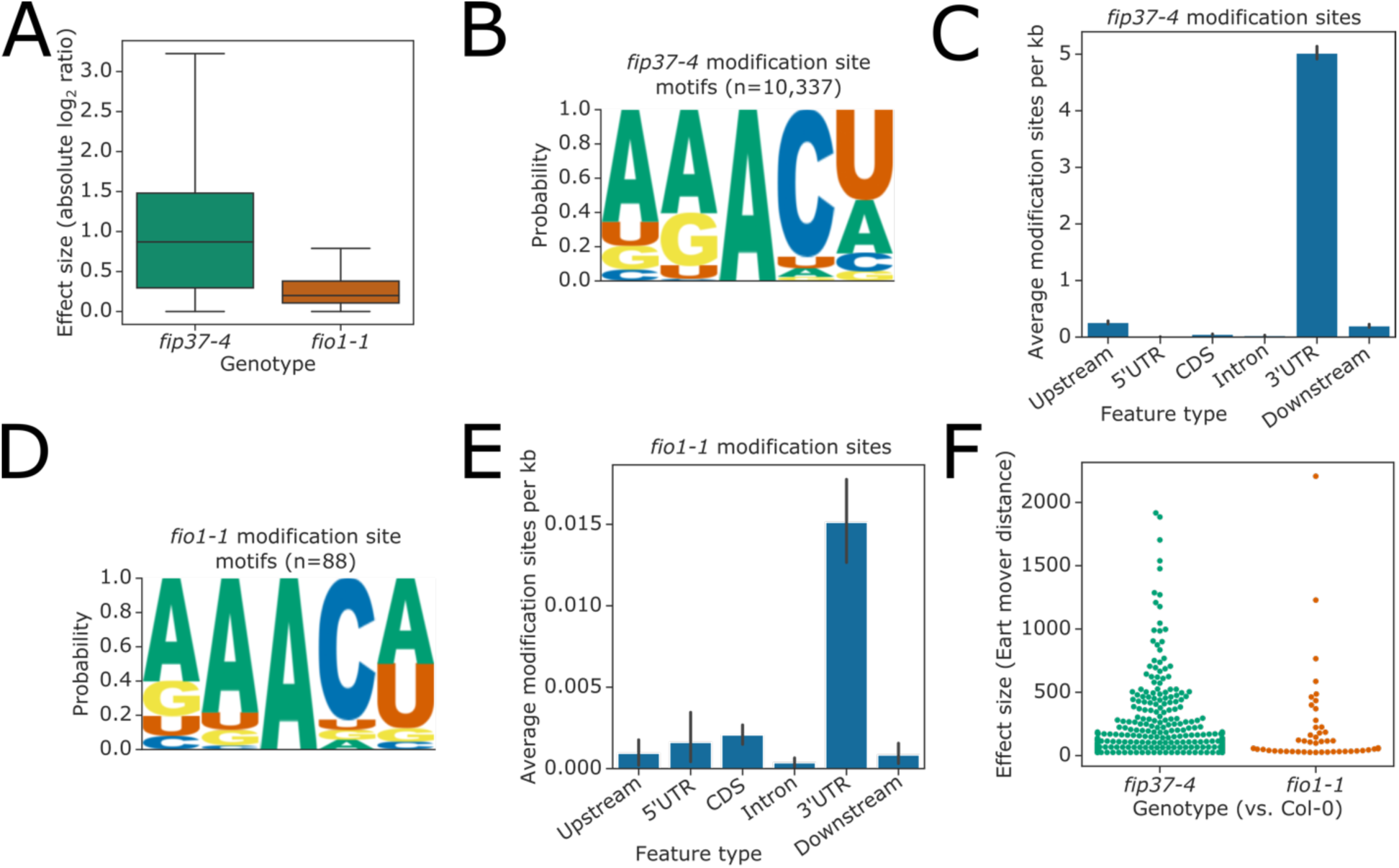
**(A)** Boxplot showing effect sizes for modification sites which have significant changes in modification rate in both *fip37-4* and *fio1-1* (intersection shown in Figure 3 -figure supplement 1A). **(B)** Sequence logo identified at FIP37-dependent modification sites detected by Yanocomp. **(C)** Bar plot showing the mean density of FIP37-dependent modification sites in different genic features annotated in Araport11. Error bars are bootstrapped 95% confidence intervals for means. **(D)** Sequence logo identified at modification sites which are significant in *fio1-1* vs Col-0 comparison, but not in *fip37-4* vs. Col-0, detected by Yanocomp, which are also supported by miCLIP. **(E)** Bar plot showing the mean density of modification sites which are significant in *fio1-1* vs Col-0 comparison, but not in *fip37-4* vs. Col-0, in different genic features annotated in Araport11. Error bars are bootstrapped 95% confidence intervals for means. **(F)** Swarmplot showing the effect size measured in Earth mover distance using d3pendr (Parker et al., 2021c) of genes with significant changes in poly(A) site choice in either *fip37-4* or *fio1-1*, compared to Col-0.

**Figure 3 – figure supplement 2.**
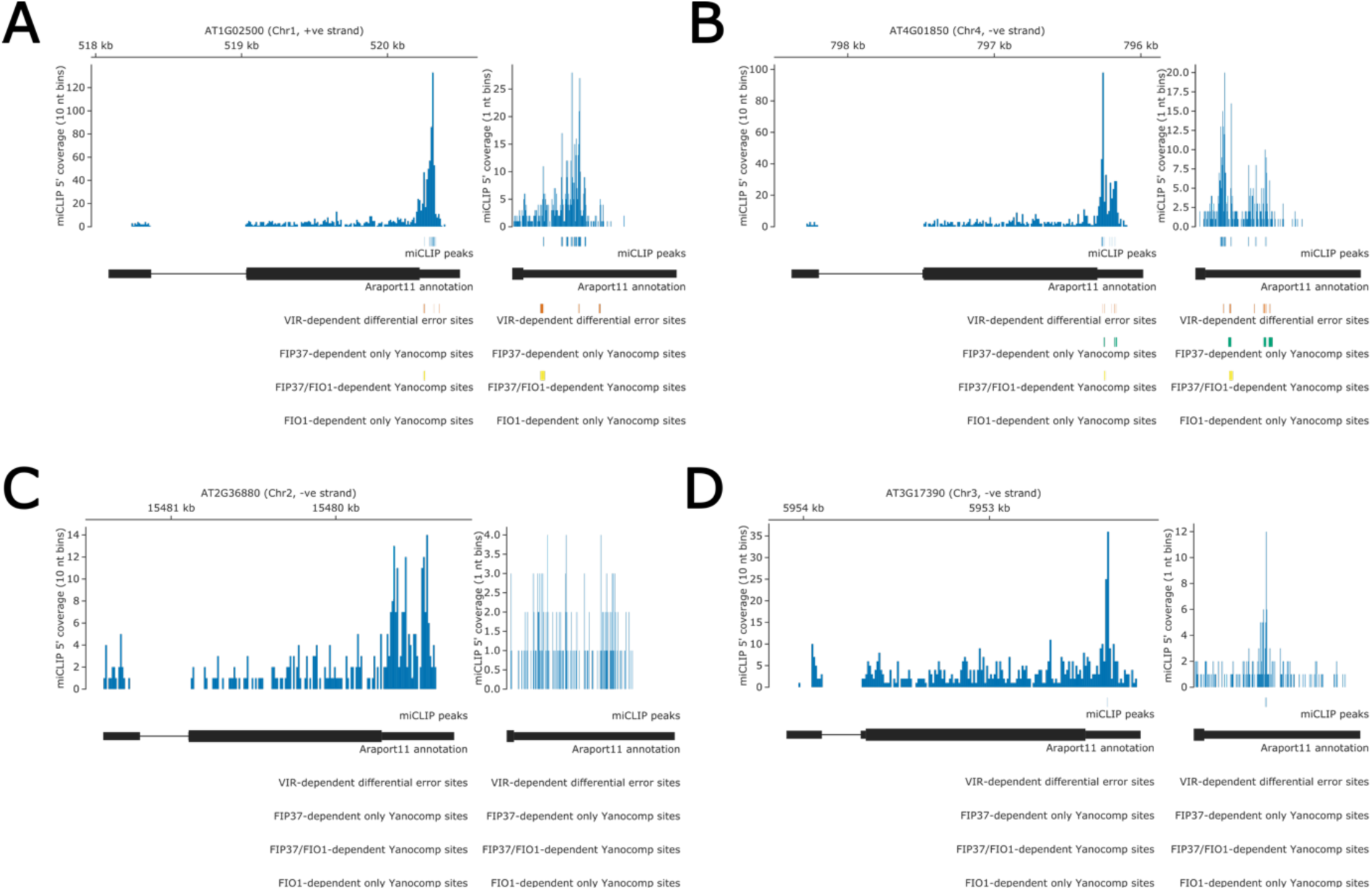
**(A-D)** Gene tracks showing predicted methylation sites in **(A)** *MAT1* **(B)** *MAT2,* **(C)** *MAT3* and **(C)** *MAT4* predicted by miCLIP (blue), differential error rate method on *vir-1* and *VIRILIZER* complemented line (orange) (Parker et al., 2020), and Yanocomp method on *fip37-4* and Col-0 lines (green). *FIP37*-dependent sites which also have significant modification rate change in *fio1-1* are shown in yellow. A magnified view of the 3’UTR is shown in the right hand panel of each gene track. No *FIO1-*dependent modification sites or gene body methylation sites were detected in *MAT1*, *MAT2, MAT3* or *MAT4*, using any of the approaches.

**Figure 3 – figure supplement 3.**
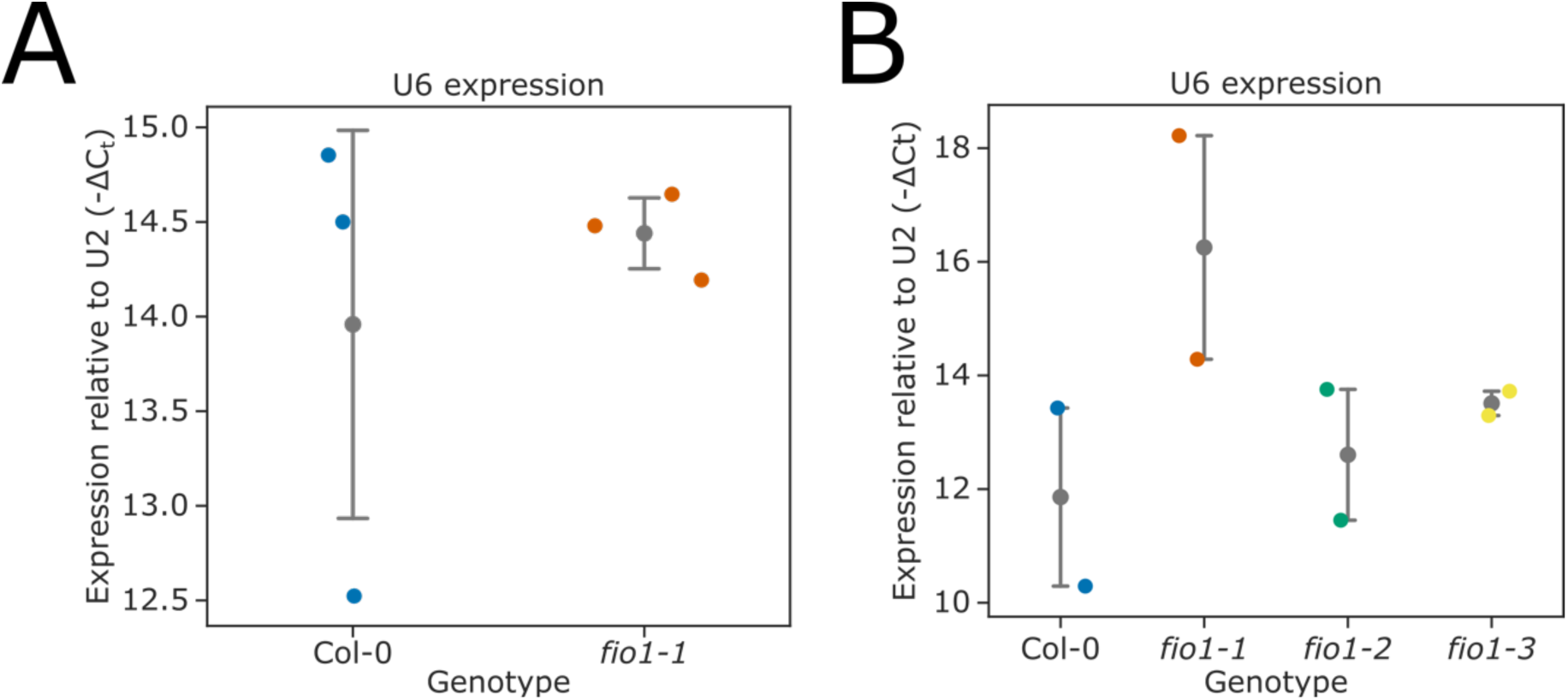
**(A-B)** Relative expression of U6 snRNA in **(A)** Col-0 and the *fio1- 1* mutant, and **(B)** Col-0, *fio1-1*, *fio1-3* and *fio1-4* mutants, measured by RT-qPCR, compared to U2 snRNA. The means of three or four technical replicates are shown for each biological replicate. Grey bars with points represent mean and 95% confidence intervals.

**Figure 4 – figure supplement 1.**
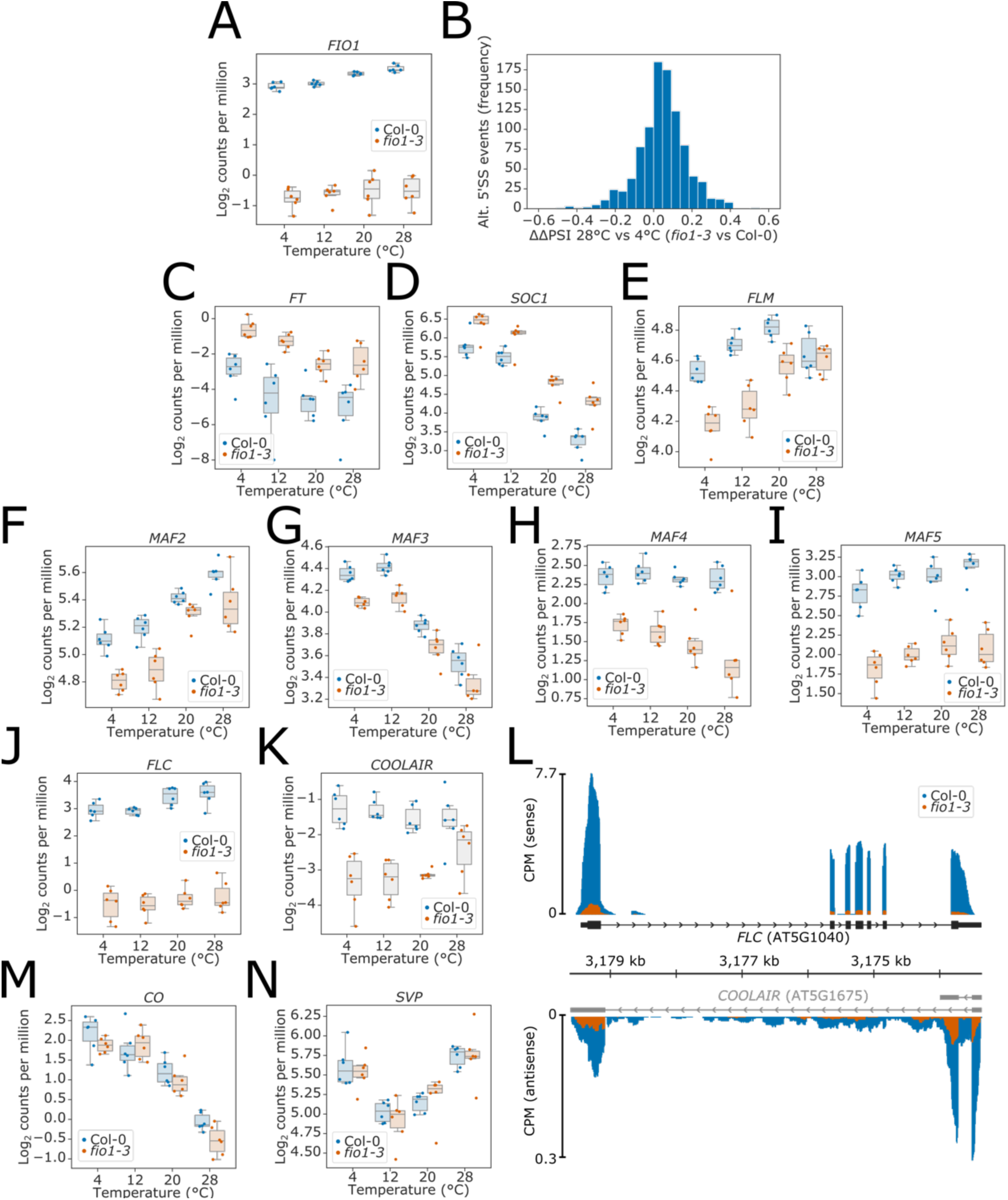
**(A)** Boxplot showing the reduction in overall gene expression of *FIO1* in *fio1-3*. **(B)** Histogram showing change in the difference between *fio1-3* and Col-0 percent splicing index (PSI) between 4°C and 28°C (ΔΔPSI). A positive ΔΔPSI indicates that there is a greater deviation from Col-0 splicing levels in *fio1-3* at 28°C than at 4°C. **(C-K, M-N)** Boxplots showing the increase in overall gene expression of the flowering time activators **(C)** *FT* (AT1G65480), **(D)** *SOC1* (AT1G65480), and reduction in the expression of the flowering repressors **(E)** *FLM* (AT1G77080), **(F)** *MAF2* (AT5G65050), **(G)** *MAF3* (AT5G65060), **(H)** *MAF4* (AT5G65070), **(I)** *MAF5* (AT5G65080), **(J)** *FLC* (AT5G10140), and **(K)** *COOLAIR* transcripts expressed antisense to *FLC.* **(L)** Gene track showing the expression of *FLC* and *COOLAIR* transcripts at 20°C in Col-0 and *fio1-3*. The mRNA-level expression of **(M)** the flowering activator *CO* (AT5G15840) and **(N)** the flowering repressor *SVP* (AT2G22540) were also not affected.

**Figure 5 – figure supplement 1.**
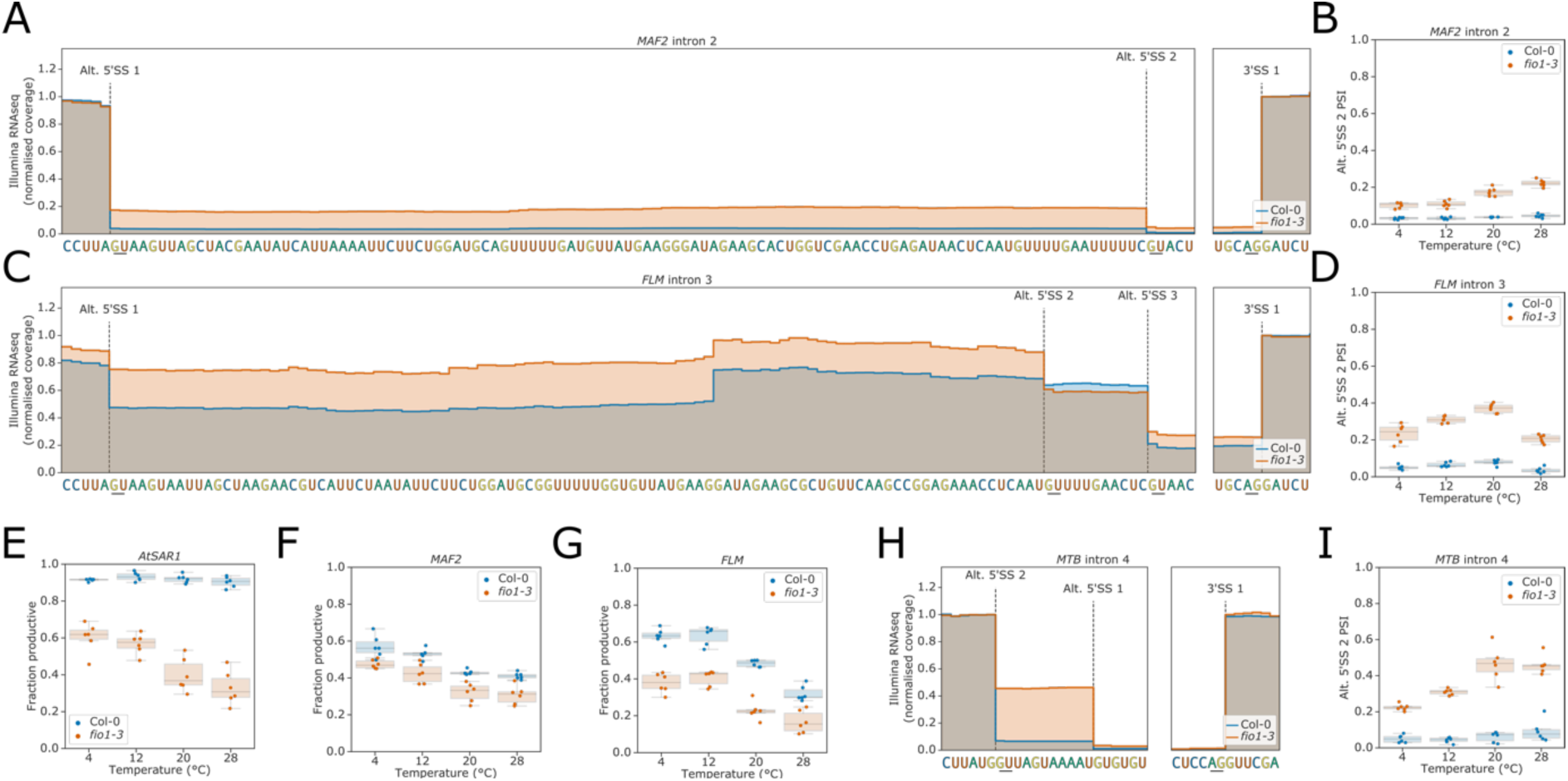
**(A)** Gene track showing alternative 5’SS usage at *MAF2* intron 2 in *fio1-3*, at 20°C. Expression is normalised by the coverage at the +1 position of the 3’SS. **(B)** Boxplot showing the change in usage of alternative 5’SS 2 of *MAF2* intron 2 at varying temperatures, in Col-0 plants and in *fio1-3*. **(C)** Gene track showing alternative 5’SS usage at *FLM* intron 3 in *fio1-3*, at 20°C. Expression is normalised by the coverage at the +1 position of the 3’SS. **(D)** Boxplot showing the change in usage of alternative 5’SS 2 of *MAF2* intron 2 at varying temperatures, in Col-0 plants and in *fio1-3*. **(E-G)** Boxplots showing the change in functional expression of **(E)** *AtSAR1,* **(F)** *MAF2* and **(G)** *FLM* in the *fio1-3* mutant, at varying temperatures, as predicted by TranSuite (Entizne et al., 2020). **(H)** Gene track showing alternative 5’SS usage at *MTB* intron 4 in *fio1-3,* at 20°C. Expression is normalised by the coverage at the +1 position of the 3’SS. **(I)** Boxplot showing the change in usage of alternative 5’SS 2 of *MTB* intron 4 at varying temperatures, in Col-0 plants and in *fio1-3*.

**Figure 5 – figure supplement 2.**
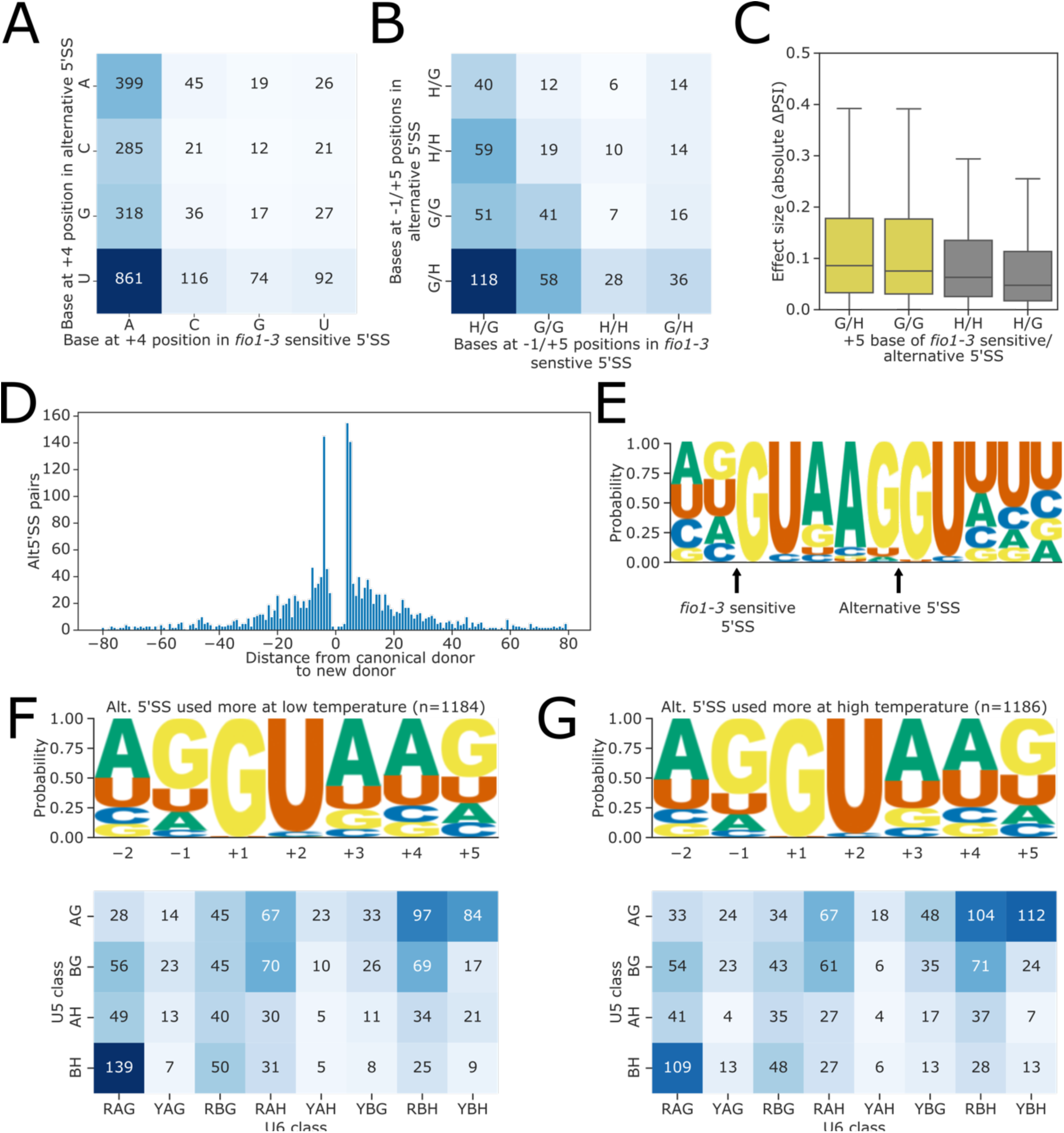
**(A)** Contingency table showing the relationship between the bases at the 4+ position for *fio1-3-*sensitive 5’SSs with reduced usage in *fio1-3* and alternative 5’SSs with increased usage in *fio1-3.* **(B)** Contingency table showing the relationship between the bases at the -1 and +5 positions for *fio1-3-*sensitive and alternative 5’SSs, where both 5’SSs in the pair have the same base at the +4 position. **(C)** Boxplot showing effect sizes of pairs of alternative 5’SSs with significantly altered usage in *fio1-3*, separated by +5 position bases (G→H indicates that 5’SS with reduced usage has G_+4_, 5’SS with increased usage has H_+4_). **(D)** Histogram showing the distance between alternative 5’SS pairs with significantly different usage in *fio1-3*. Negative distances represent shifts towards greater usage of upstream 5’SSs, whilst positive distances represent shifts towards greater usage of downstream 5’SSs. **(E)** Sequence logo for alternative 5’SS pairs where the distance between the *fio1-3* sensitive 5’SS and the alternative 5’SS is exactly +5 nt. **(F-G)** Sequence logos for 5’SSs which have **(F)** increased and **(G)** decreased retention in the *fio1-3* mutant. Motifs are shown for the —2 to +5 positions of the 5’SS.

**Figure 6 – figure supplement 1.**
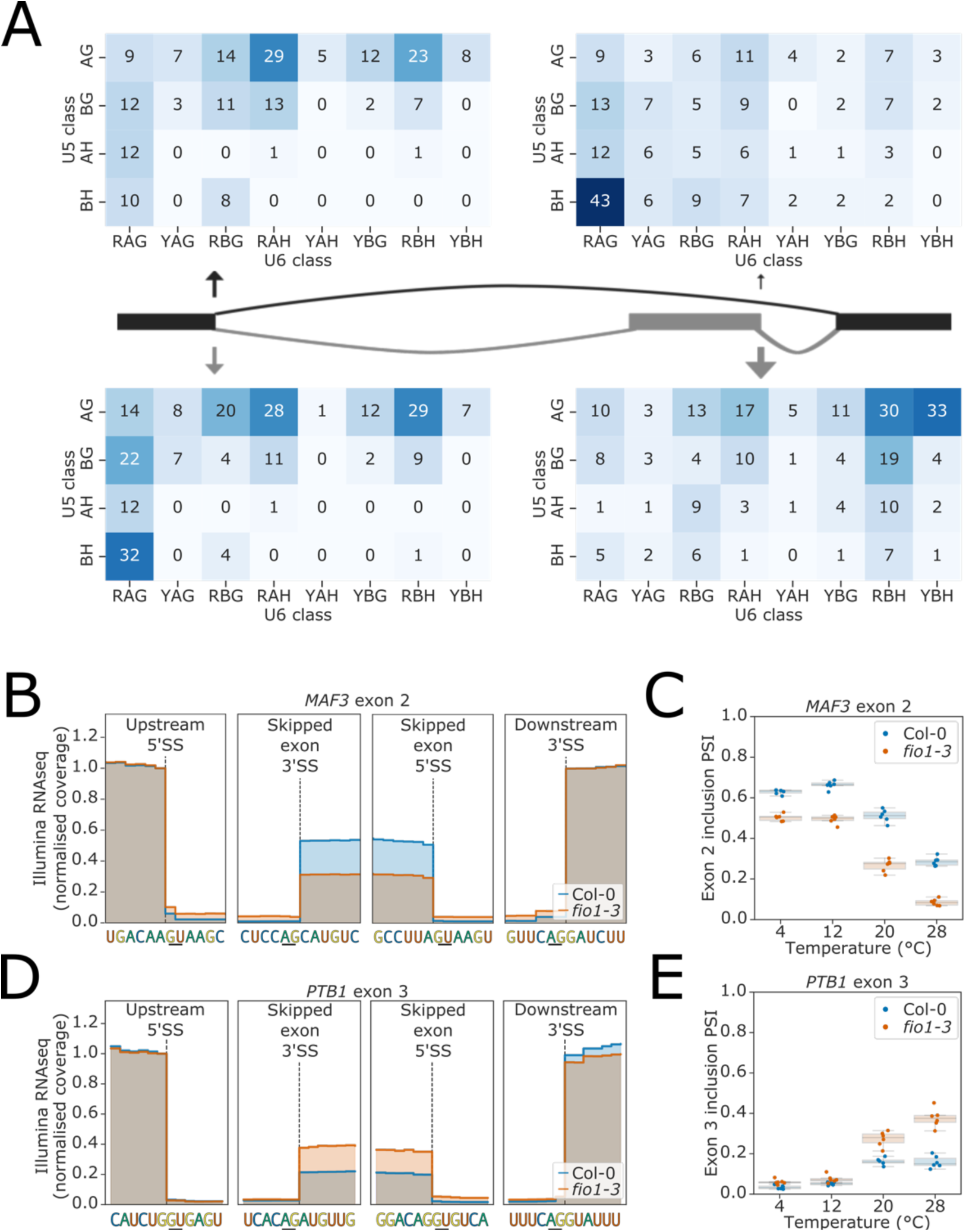
**(A)** Heatmaps showing the distribution of U5 snRNA and U6 snRNA interacting sequence classes for 5’SSs of exons (right) and upstream 5’SSs (left) which have increased (above) and decreased (below) skipping in *fio1-3*. U5 classes are based upon the distance of the —2 to —1 positions of the 5’SS from the consensus motif AG. U6 classes are based upon the distance of the +3 to +5 positions of the 5’SS from the consensus motif RAG. **(B)** Gene track showing change in exon skipping at *MAF3* exon 2 in *fio1-3*, at 20°C. Expression is normalised by the coverage at the +1 position of the upstream 5’SS. **(B)** Boxplot showing the change in *MAF2* exon 2 inclusion at varying temperatures, in Col-0 and in *fio1-3*. **(B)** Gene track showing change in exon skipping at *PTB1* exon 3 in *fio1-3*, at 20°C. Expression is normalised by the coverage at the +1 position of the upstream 5’SS. **(B)** Boxplot showing the change in *PTB1* exon 3 inclusion at varying temperatures, in Col-0 and in *fio1-3*.

**Figure 7 – figure supplement 1.**
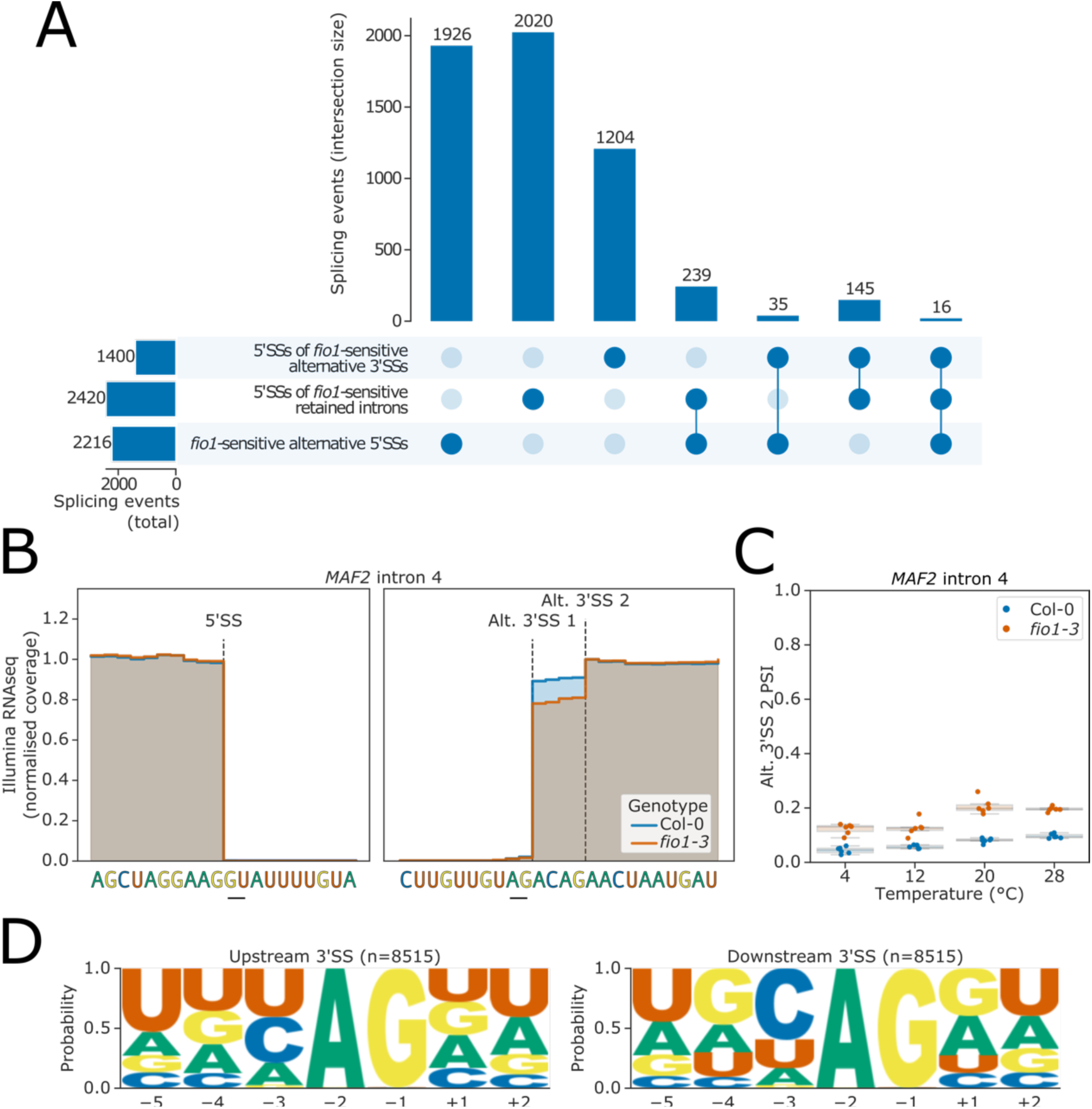
**(A)** Upsetplot showing the overlap of *fio1-*sensitive 5’SSs involved in alternative 5’SS usage, with 5’SSs at *fio1-*sensitive intron retention events and alternative 3’SSs. **(B)** Gene track showing alternative 3’SS usage at *MAF2* intron 4 in *fio1-3*, at 20°C. Expression is normalised by the coverage at the −1 position of the 5’SS. **(C)** Boxplot showing the change in usage of alternative 3’SS 2 of *MAF2* intron 4 at varying temperatures, in Col-0 and *fio1-3*. **(D)** Sequence logos for all upstream and downstream alternative 3’SS pairs identified from the RNA-Seq data.

**Figure 8 – figure supplement 1.**
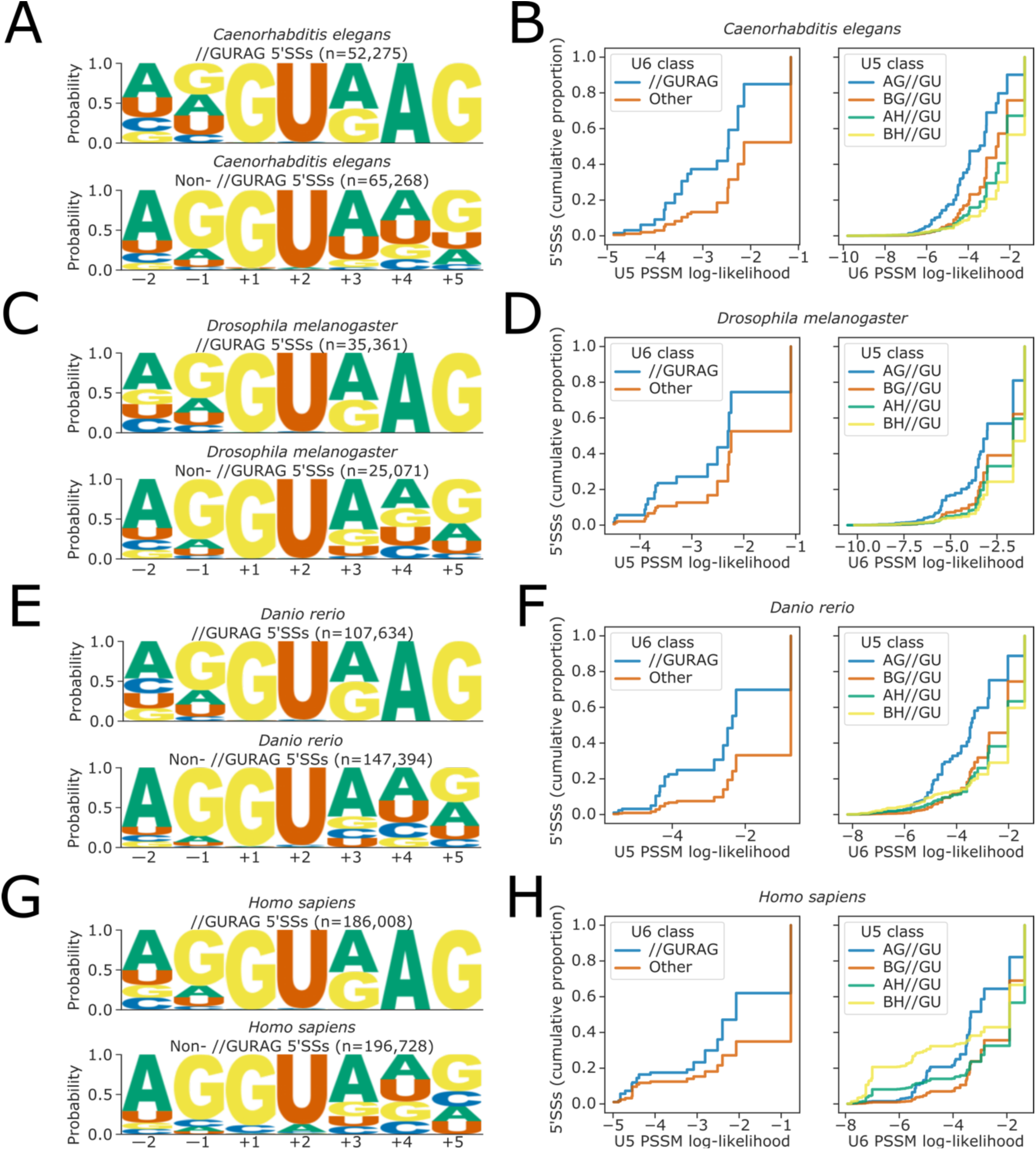
**(A, C, E, G)** Sequence logos showing base frequency probabilities at —2 to +5 positions for (above) 5’SSs with //GURAG sequence and (below) all other 5’SSs, in the organisms **(A)** *C. elegans,* **(C)** *D. melanogaster,* **(E)** *D. rerio* and **(G)** *H. sapiens*. **(B, D, F, H)** Empirical cumulative distribution function of U5 snRNA (left panels) and U6 snRNA (right panels) PSSM log-likelihood scores for 5’SSs falling into different U6 (left panels) or U5 (right panels) sequence classes, for all 5’SSs in the organisms **(B)** *C. elegans,* **(D)** *D. melanogaster,* **(F)** *D. rerio* and **(H)** *H. sapiens*. PSSM scores are calculated using a PSSM derived from all 5’SSs in the corresponding annotation of each organism, at the —2 to —1 positions of the 5’SS for U5 and the +3 to +5 positions for U6, inclusive.

**Figure 8 – figure supplement 2.**
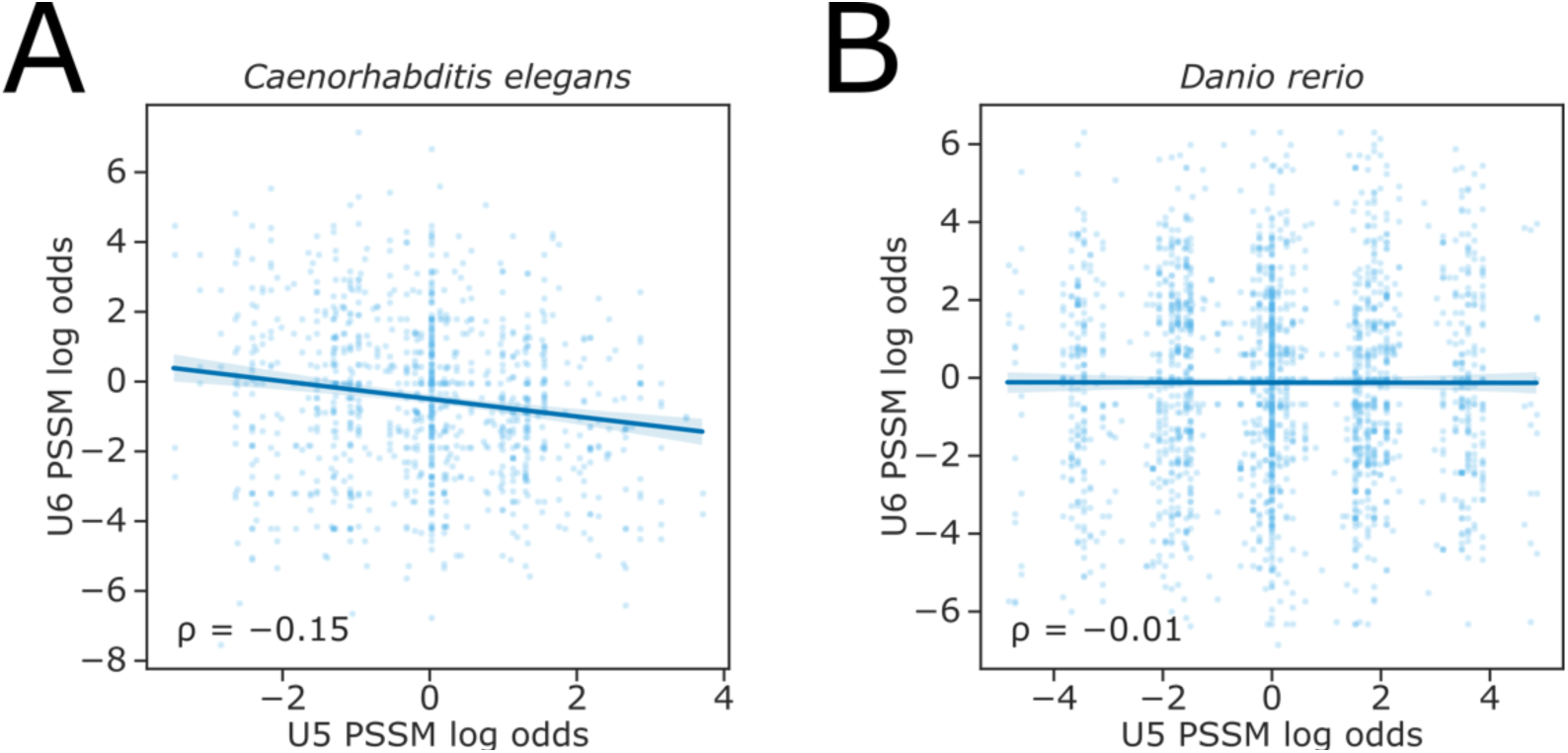
**(A-B)** Scatterplot showing the ratio of PSSM log-likelihoods (log-odds ratio) for U5 snRNA and U6 snRNA interacting sequences, at pairs of upstream and downstream alternative 5’SSs in the **(A)** *C. elegans* or **(B)** *D. rerio* reference annotation. A positive log-odds ratio indicates that the PSSM score of the upstream 5’SS is greater than that of the downstream 5’SS.

**Figure 9 – figure supplement 1.**
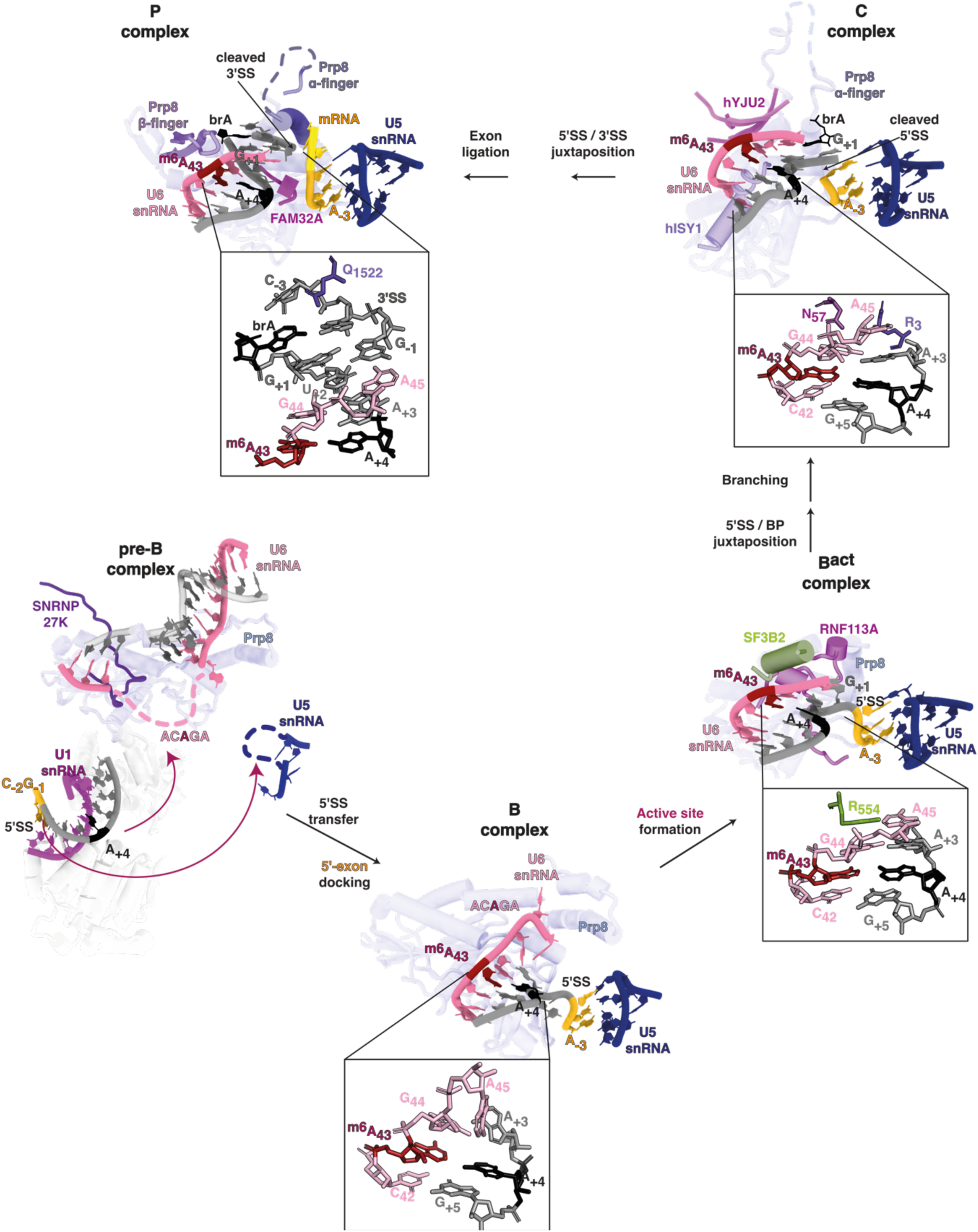
Tracing m^6^A modified U6 snRNA in cryo-EM structures of different spliceosomal complexes with cryo-EM reveals in pre-B complex (PDB 6QX9, Charenton et al., 2019), the ACAGA box is flexible and disordered. In B complex (PDB 6AHD, Bertram et al., 2017), m^6^A forms a trans Hoogsteen sugar edge interaction with 5’SS A_+4,_ capping the U6/5’SS helix by stacking. A kink in the 5’SS is detectable and U5 snRNA loop 1 (dark blue) is docked on the upstream exon (yellow). In the B^act^ complex (PDB 5Z56, Zhang et al., 2018), U6 G44 pairs with 5’SS A_+3,_ capping and stabilising the U6/5’SS helix. This interaction is further stabilised by R554 of the spliceosomal protein SF3B2. Consequently, the role of m^6^A:5’SS A_+4_ in stabilising the helix appears less important at this stage. In C complex (PDB 6ZYM, Bertram et al., 2020), the 5’SS/U6 helix is stabilised by R3 of Prp8 and N57 of hYJU2. Rearrangements in P complex for the second splicing reaction suggest the stabilising role of U6 m^6^A43:5’SS A_+4_ becomes important again because 5’SS A_+3_ pivots away to form new stacking interactions (PDB 6QDV, Fica et al., 2019). The 3’SS G_+1_ stacks on U6 snRNA A45, which now forms a non-canonical base-pair with 5’SS U_+2_. A continuous helix stack involving the U6/5’SS interaction between U6 m^6^A43 and 5’SS A_+4,_ and the docked 3’SS, is capped by interaction between 3’SS_-3_ and Prp8 Q1522. A domain of the major architectural spliceosomal protein, Prp8, is depicted in the background as a constant reference through each stage. The methyl group of U6 snRNA m^6^A43 is not modelled in the structures due to lack of resolution.

## LIST OF SUPPLEMENTARY FILES

Supplementary file 1: Read statistics for Nanopore DRS and Illumina RNAseq datasets.

Supplementary file 2: Oligos and primers used in this study.

## LIST OF SOURCE DATASETS

Figure 2 source data 1: Mutations identified in early flowering and *MAF2* splicing screen, detected by artMap.

Figure 2 source data 2: Sanger sequencing products for *FIO1* cDNAs in *fio1-1* and *fio1-4* mutant.

Figure 2 source data 3: Sanger sequencing product alignments for *FIO1* cDNAs in *fio1-1* and *fio1-4* mutant.

Figure 2 source data 4: MAF2 intron 3 splicing qPCR data for Col-0, *fio1-1* and *fio1-3* mutants.

Figure 2 source data 5: Flowering time data for *fio1-1*, *fio1-3* and *maf2* mutants.

Figure 3 source data 1: Differential modification sites detected from a three-way comparison of Col-0, *fip37-4* and *fio1-1* mutants using nanopore DRS data.

Figure 3 source data 2: Differential poly(A) site usage results for Col-0 vs. *fip37-4* identified using nanopore DRS data.

Figure 3 source data 3: Differential poly(A) site usage results for Col-0 vs. *fio1-1* identified using nanopore DRS data.

Figure 3 source data 4: m^6^A-IP qPCR data for U6 and U2 snRNAs in Col-0 and *fio1-1, fio1-3 and fio1-4* mutants.

Figure 4 source data 1: Differential gene expression analysis results from Illumina RNA-Seq experiment on Col-0 and *fio1-3* mutants at four temperatures.

Figure 4 source data 2: *De novo* transcriptome assembly generated from Col-0 and *fio1* mutant nanopore DRS and Illumina RNA-Seq

Figure 4 source data 3: Differential splicing analysis results from Illumina RNA-Seq experiment on Col-0 and *fio1-3* mutants at four temperatures.

Figure 5 source data 1: Differential productive transcription/NMD analysis results from Illumina RNA-Seq experiment on Col-0 and *fio1-3* mutants at four temperatures.

